# *p*-Brain: An Automated MRI Pipeline for Cerebral Perfusion, Microvasculature, and Blood–Brain Barrier Permeability Estimation

**DOI:** 10.64898/2026.02.15.705995

**Authors:** Edis D. Tireli, Stig P. Cramer, Ulrich Lindberg, Derya Tireli, Mark B. Vestergaard, Henrik B. W. Larsson

## Abstract

We present *p*-Brain, an end-to-end neuroimaging analysis framework for reproducible, automated quantitative DCE-MRI analysis at scale. From standard acquisitions, *p*-Brain estimates baseline relaxation parameters, converts signal to gadolinium concentration, derives arterial and venous input functions using convolutional neural network (CNN) slice selection and ROI segmentation, and produces voxelwise maps with regional and whole-brain summaries. The pipeline implements Patlak graphical analysis to estimate the blood–brain barrier influx constant (*K*_*i*_) and plasma volume fraction (*v*_*p*_), and performs model-free residue deconvolution with Tikhonov regularisation to estimate cerebral blood flow (CBF), mean transit time (MTT), and capillary transit-time heterogeneity (CTH) from the same DCE dataset. *p*-Brain exports analysis-ready outputs, intermediate readouts, structured runtime metadata, and stage-level quality control artifacts to support auditability in batch processing. We evaluate the framework on a technically uniform set of 97 DCE-MRI scans from 58 healthy human participants, and show close agreement between automated Patlak *K*_*i*_ summaries and an established reference workflow. A companion macOS desktop application supports batch execution, job monitoring, and rapid review of curves and maps. *p*-Brain is open-source and configurable, enabling extension to additional kinetic models.

## 1 Introduction

The blood–brain barrier (BBB) regulates cerebral homeostasis, and BBB dysfunction is implicated in many neurological disorders. Accurate quantification of BBB permeability can be used to study neurological diseases such as multiple sclerosis, brain tumors, and neurodegenerative disorders (Cramer et al., 2014; Okuchi et al., 2019; Heye et al., 2014; Chagnot et al., 2021; Raja et al., 2018). Dynamic contrast-enhanced MRI (DCE-MRI) enables in vivo quantification of subtle BBB leakage (Varatharaj et al., 2019; Ingrisch et al., 2012) via tracer-kinetic models such as Patlak graphical analysis (Patlak et al., 1983) and other compartmental models. These approaches estimate permeability-related parameters (e.g., *K*_*i*_) and complementary metrics. Prior work supports the biological validity and clinical relevance of these measurements: DCE-MRI-derived *K*_*i*_ shows expected gray/white matter differences and lesion sensitivity (Varatharaj et al., 2019), can be reproducible even at low leakage levels (Cramer et al., 2023a), and has demonstrated prognostic value (e.g., optic neuritis to multiple sclerosis conversion) (Cramer et al., 2015). Beyond demyelinating disease, BBB permeability mapping has been applied in acute ischemic stroke and reperfusion injury (Villringer et al., 2017; Arba et al., 2021), traumatic brain injury (Ware et al., 2022), seizure disorders such as new-onset refractory status epilepticus (Li et al., 2023), autoimmune and systemic inflammatory disease including autoimmune encephalitis and systemic lupus erythematosus (Ji et al., 2024; Chi et al., 2019), infectious neuroinflammation such as neuroborreliosis (Lindland et al., 2023), and HIV-associated neurocognitive impairment (Chaganti et al., 2019). Together, these findings motivate automated, user-independent tools for estimating BBB permeability and related perfusion parameters at scale.

Despite the potential of DCE-MRI, analysis often remains labor-intensive. Manual ROI definition (e.g., arterial input functions and tissue curves) and supervision of model fitting introduce operator variability and limit scalability (Varatharaj et al., 2024a; Beresford et al., 2006; Calamante, 2013; Keil et al., 2017). This motivates fully automated pipelines that integrate preprocessing, ROI extraction, and pharmacokinetic modelling while preserving quantitative fidelity (Varatharaj et al., 2024a; Barnes et al., 2015; Nalepa et al., 2020).

To overcome these challenges, we introduce *p*-Brain, a fully automated pipeline that integrates convolutional neural network (CNN)-driven ROI segmentation, adaptive automated arterial input extraction, and tracer-kinetic modeling (Patlak graphical analysis) alongside model-free deconvolution for perfusion and microvascular transit metrics. The design targets complete removal of user interaction while preserving agreement with established analyses. In addition to BBB permeability and plasma volume, *p*-Brain estimates cerebral blood flow (CBF) and residue-derived descriptors of mean transit time (MTT) and capillary transit-time heterogeneity (CTH), enabling interrogation of both bulk perfusion and microvascular dispersion within the same workflow. The framework is designed to be extensible, enabling additional kinetic models to be incorporated as needed. Our pipeline delivers robust, reproducible metrics at multiple analytical scales (voxel-wise, regional, whole-brain) without requiring user interaction.

The workflow is designed to be configurable, with key analysis choices (e.g., input-function strategy, curve extraction, and optional normalization) documented online.

The emergence of fast, deep learning-based neuroimaging pipelines (e.g., FastSurfer (Henschel et al., 2020)) enables whole-brain structural segmentation within minutes, offering a potential solution for consistent ROI definition across subjects. *p*-Brain leverages these advances to automatically generate tissue masks and parcellations for regional summarisation, allowing however the user to use their own choice of segmentation software as well. In this paper, we describe the implementation of *p*-Brain and validate its performance against reference methods and literature values.

Data from DCE-MRI are obtained according to clinical protocols but at minimum requires a 3D *T*_1_ weighted image, a series of inversion times, *T*_*I*_, recovery sequences for the estimation of *T*_1_ and *M*_0_ parameters, as well as the DCE-MR image. Temporal resolution and duration should be optimized to ensure precise kinetic modeling. MRI data undergo axial reslicing of structural volumes to the native DCE frame.

## 2 Results

We applied *p*-Brain to a technically uniform subset of 97 DCE-MRI scans from 58 healthy participants with no macroscopic lesions on structural MRI (Methods). Each dataset was processed end-to-end with identical settings in fully automatic mode. Deliverables include voxelwise, regional, and whole-brain outputs of BBB influx (*K*_*i*_), plasma volume (*v*_*p*_), cerebral blood flow (CBF), capillary transit-time heterogeneity (CTH), and mean transit time (MTT), and structured run summaries saved alongside the maps.

Figures 1–9 show representative outputs from a fully automatic run. Voxelwise maps capture within-scan spatial heterogeneity, while parcel- and cohort-level summaries support group comparisons and downstream statistics.

**Figure 1:**
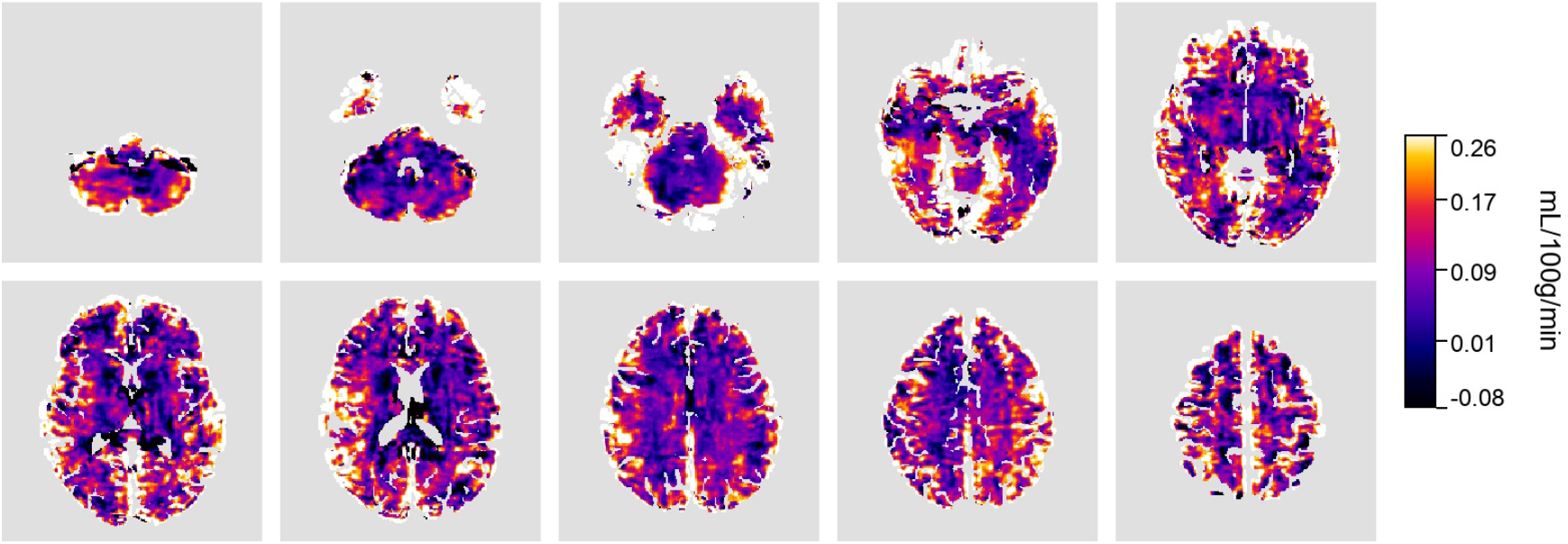
Automated voxelwise *K*_*i*_ map in units of mL*/*100g*/*min of the representative example. Local heterogeneities can be seen within and across tissue.

### Voxelwise maps

Voxelwise maps quantify model-derived parameters at the native spatial scale, preserving continuous structure and enabling localized assessment of heterogeneity that can be obscured by regional aggregation.

To illustrate the voxelwise approach, we turn first to *K*_*i*_, a parameter by default within the pipeline estimated from Patlak graphical analysis (see Patlak graphical analysis) describing the unidirectional transfer of contrast agent from plasma to tissue, reflecting the blood–brain barrier permeability. The map in Figure 1 displays the voxelwise estimation of this parameter across the entire brain volume, providing a spatially continuous quantification of barrier function.

On maps such as these, focal or asymmetric *K*_*i*_ elevations could signify localised blood–brain barrier leakage or microvascular dysfunction, which may be relevant in both diffuse and focal pathologies. Turning to perfusion, Figure 2 shows voxelwise cerebral blood flow (CBF) estimated by model-free residue deconvolution (elaborated in Modelling).

**Figure 2:**
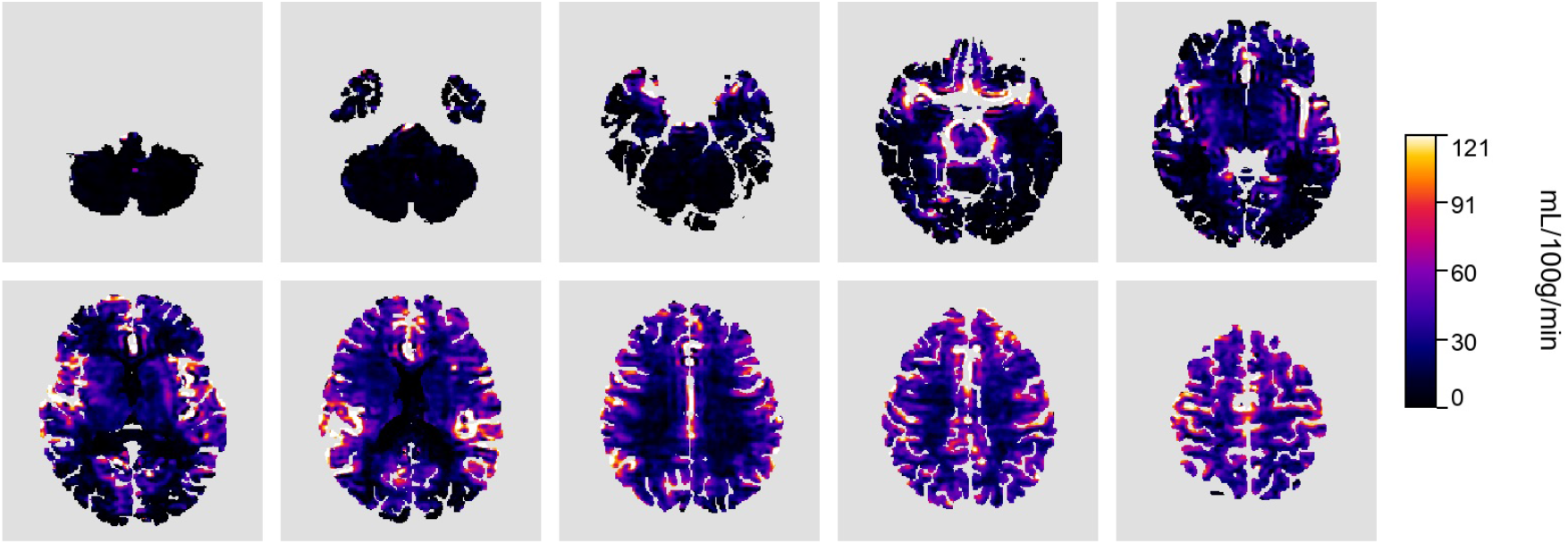
Voxelwise CBF from model-free deconvolution of the representative participant in units of mL*/*100g*/*min. Local heterogeneities can be seen as well as elevated CBF around the neurovasculature; notably in this dataset, the arteries forming the circle of Willis are visible in the lower slices.

The voxelwise plasma volume map (Figure 3) reports the fractional intravascular volume *v*_*p*_ estimated from the Patlak intercept. In practice, *v*_*p*_ highlights regions where the tissue signal is dominated by blood rather than extravascular uptake, and it is therefore sensitive to vascular density, large-vessel partial-volume effects, and residual vascular signal in the tissue curves. Interpreting *v*_*p*_ alongside *K*_*i*_ helps separate true exchange from apparent leakage driven by vascular volume (e.g., high *v*_*p*_ with low *K*_*i*_ vs. low *v*_*p*_ with elevated *K*_*i*_), and provides a complementary context for perfusion measures such as CBF.

**Figure 3:**
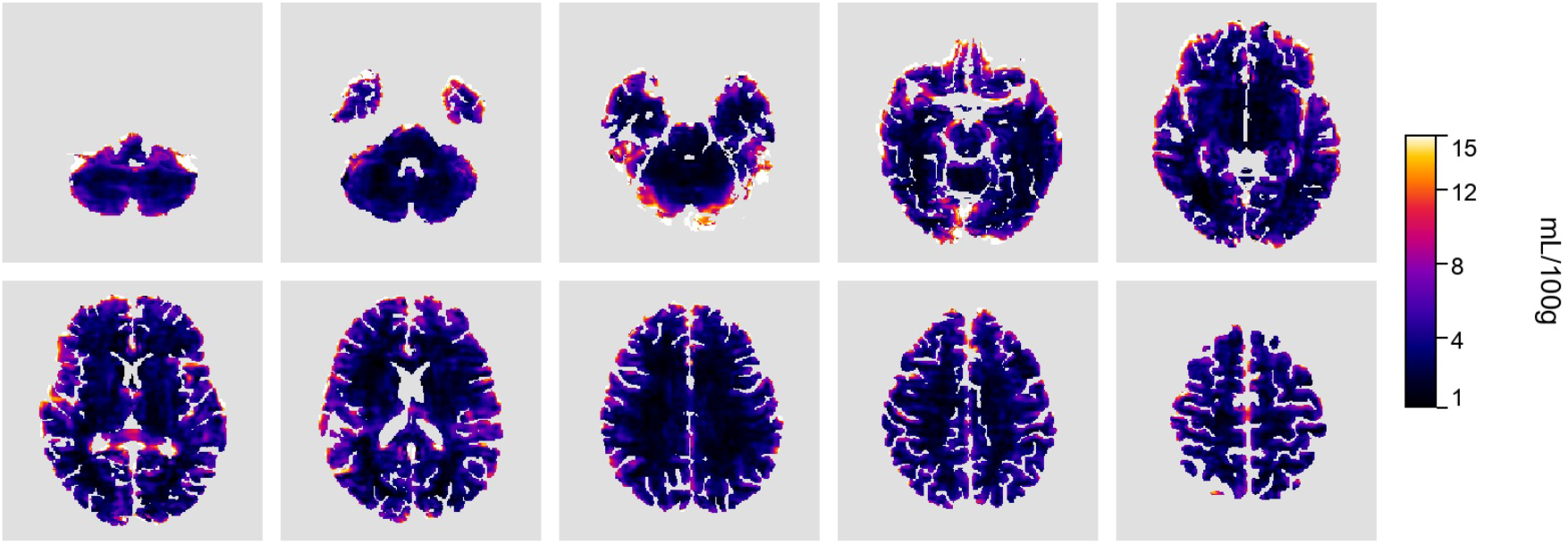
Voxelwise plasma volume (*v*_*p*_; Patlak intercept) for the representative dataset in units of mL*/*100g.

Capillary transit-time heterogeneity (CTH) summarises the dispersion of microvascular transit times estimated from the residue/outflow function (Angleys et al., 2015), shown in Figure 4. In practice, elevated CTH reflects a broader distribution of capillary passage times and can indicate increased microvascular flow heterogeneity, which may reduce effective oxygen extraction even when bulk flow is preserved (Larsson et al., 2017a; Vestergaard et al., 2023a). CTH maps can therefore be relevant when probing microvascular dysfunction in settings such as small-vessel disease, inflammatory or metabolic stress, and peri-lesional hemodynamic remodeling, and they are most interpretable when considered jointly with CBF and MTT (e.g., heterogeneous vs. uniformly delayed transit).

**Figure 4:**
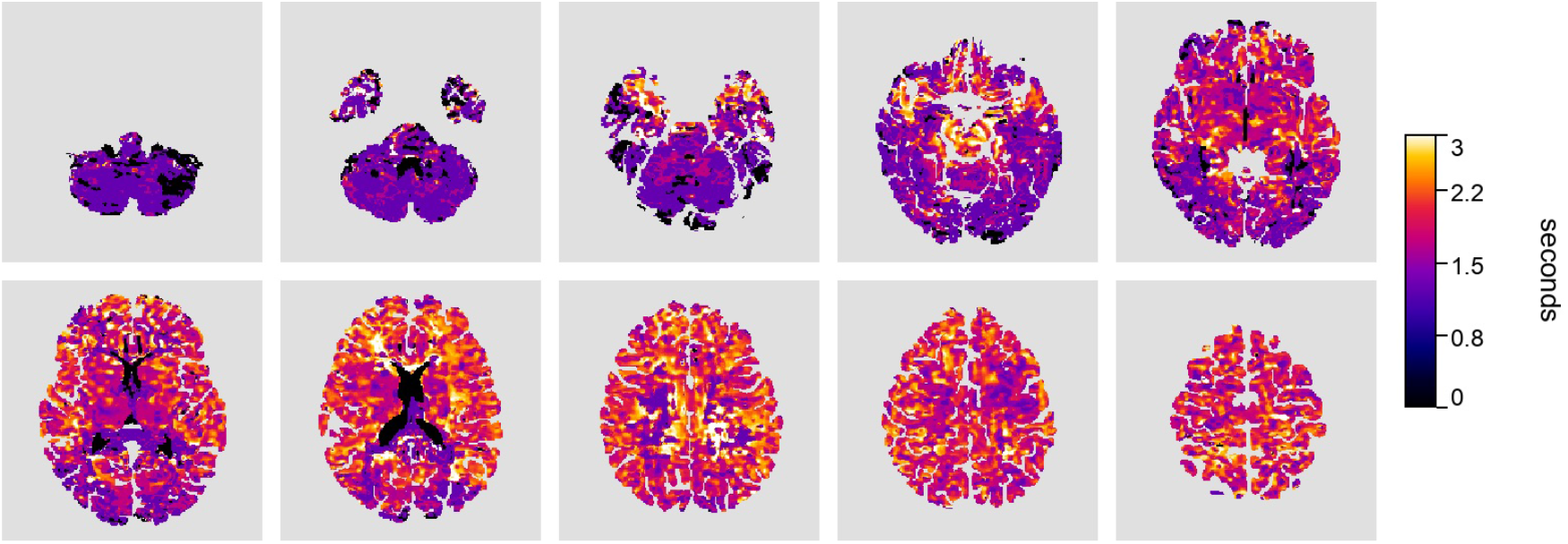
Voxelwise CTH derived from the normalized outflow *h*(*t*) = −*r*^*′*^(*t*)*/*∫(−*r*^*′*^) with units in s.

Fine-scale spatial texture in voxelwise CTH can reflect physiologic heterogeneity but is also sensitive to noise and deconvolution regularization; *p*-Brain therefore exports voxelwise maps alongside parcel and cohort summaries to support reproducible interpretation across subjects.

Mean transit time (MTT) summarizes the average duration of contrast passage through the microvasculature as derived from the first moment of the residue function (Figure 5). Elevated MTT indicates delayed transit and may be observed with reduced perfusion pressure, increased vascular resistance, or reliance on collateral pathways, whereas lower MTT is consistent with faster passage. MTT can be useful for characterizing perfusion delay in cerebrovascular disease and for contextualizing permeability or tissue injury measures; interpreted alongside CBF and CTH it helps distinguish slow-but-uniform flow from heterogeneous microvascular transit.

**Figure 5:**
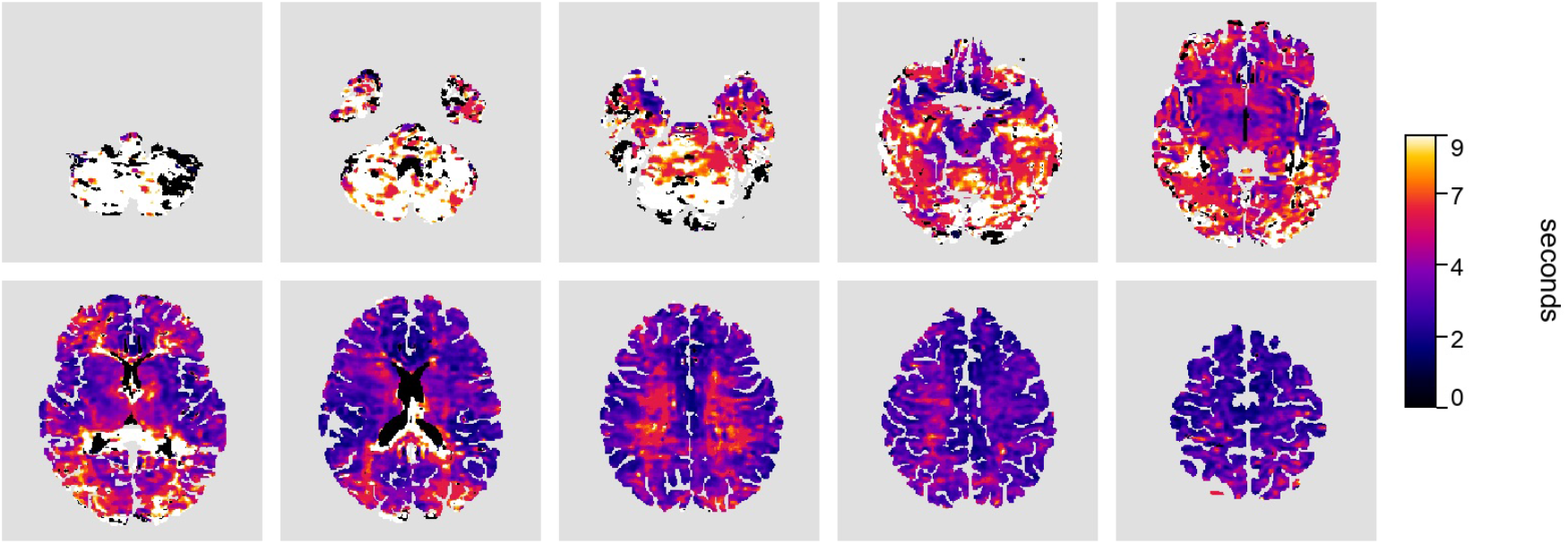
Voxelwise MTT derived from the first moment of the residue function with units in s.

### Regional and parcellated organization

Voxelwise maps preserve fine spatial detail, but they are cumbersome for cohort statistics and can be disproportionately influenced by local noise, partial volume, and small residual misregistrations. If a high resolution 3D T1-weighted scan is available, *p*-Brain converts every quantitative output into a common anatomical representation by propagating the parcellation from segmentation^1^ into DCE space and summarising each map by parcel; the mean or median value (user configured) is found within the ROI of the parcel. This produces, for every scan, a standardised regional feature set that can be analysed directly (e.g., longitudinal tracking, group comparisons, and multi-site harmonisation) without additional scripting.

Figure 6 shows parcel-level CBF, illustrating how the workflow turns a voxelwise perfusion field into a regional map that is immediately comparable across participants. Figure 7 shows the corresponding parcel-wise Patlak *K*_*i*_, providing an interpretable regional summary of BBB leakage that supports localisation of effects and downstream statistics. Additional parcel-level maps (CTH, MTT, and *v*_*p*_) are provided in the Appendix (Figures 25–27).

**Figure 6:**
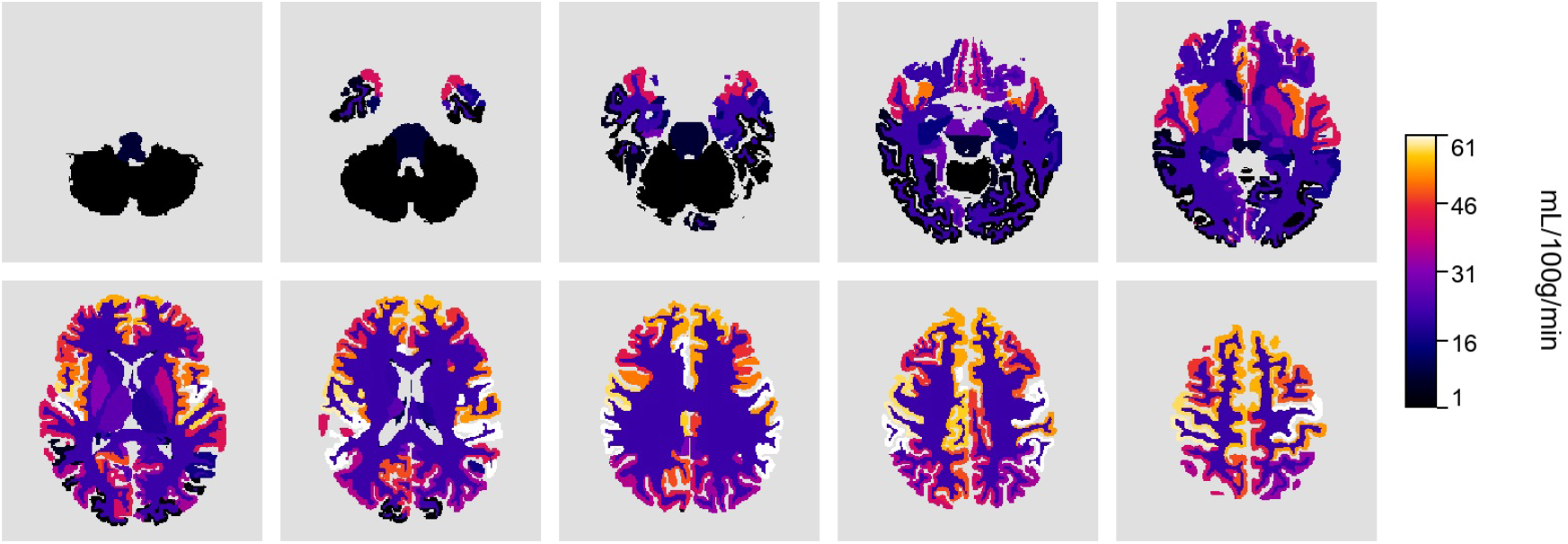
Parcel-level CBF in the representative participant. Units: mL*/*100g*/*min.

**Figure 7:**
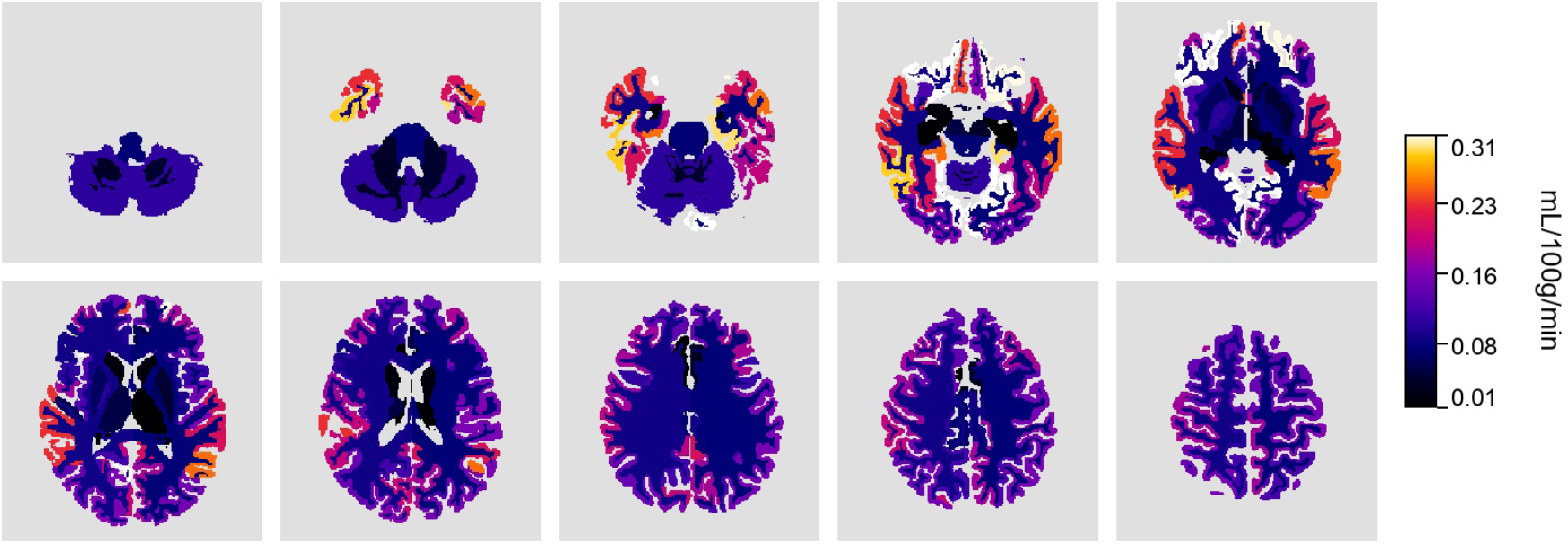
Parcel-level *K*_*i*_ within the representative participant.

These regional exports are therefore more than visual aids: they constitute ready-made datasets that can be pulled directly into statistical software, shared across centres, or revisited as new hypotheses emerge. By automating the parcellation, *p*-Brain lets researchers focus on biological interpretation—mapping where perfusion, permeability, and microvascular timing diverge—rather than on the logistics of producing the maps.

### Cohort-level atlas projection

Group-level summaries are essential for quantitative DCE-MRI research because they enable comparisons against control populations, between subgroups within a disease, and across disease entities or phenotypes. *p*-Brain automatically aggregates parcel-level summaries across subjects and projects cohort means onto a common reference segmentation to provide cohort-level context for permeability and perfusion metrics. Figure 8 shows the cohort-mean Patlak *K*_*i*_ projected onto the reference atlas. Additional cohort-mean atlas projections (CBF, CTH, and MTT) are provided in the Appendix (Figures 22–24).

**Figure 8:**
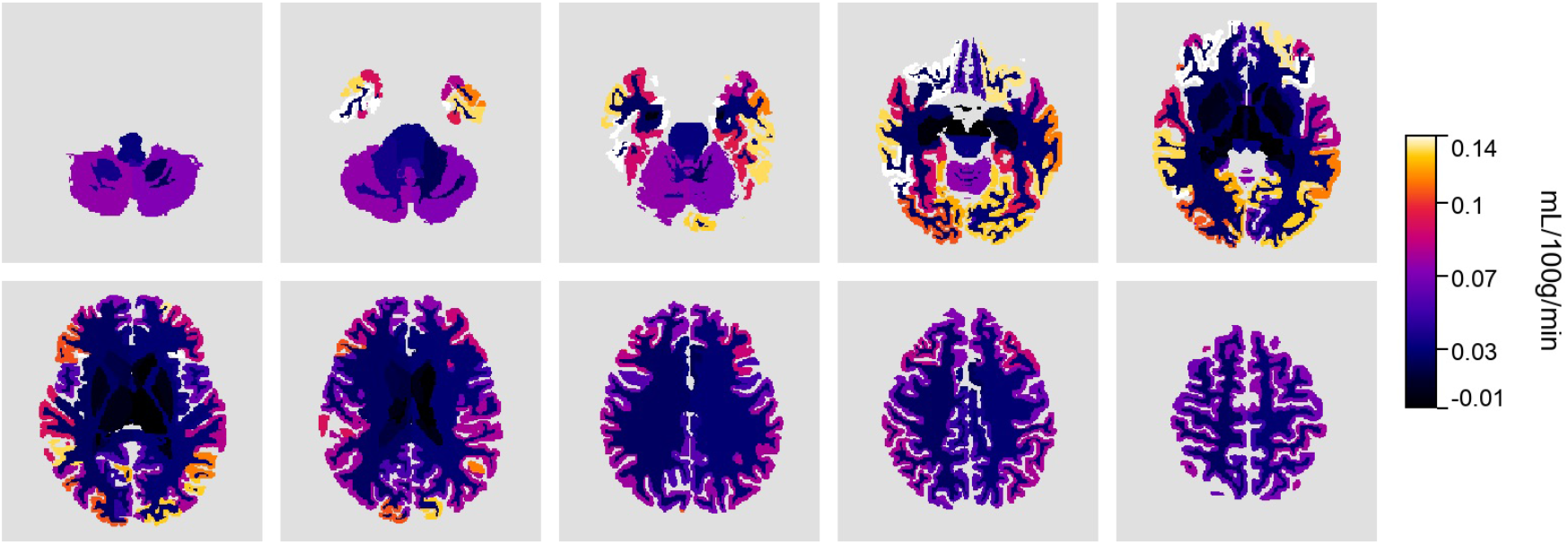
Cohort-mean *K*_*i*_ projected onto a reference atlas. Units: mL*/*100g*/*min.

Summarising these maps by anatomical parcels reduces the influence of voxelwise noise and small registration errors, emphasises regionally coherent effects, and enables consistent group-level comparisons.

### Whole-brain medians

For scenarios requiring concise descriptors, *p*-Brain reports tissue-specific whole-brain medians derived from robust aggregate curves. Figure 9 shows that these medians preserve the gray/white ordering while condensing each scan to a handful of values. Such summaries can populate electronic health records, feed downstream statistical models, or serve as quality-control references alongside the richer voxelwise and parcelwise outputs.

**Figure 9:**
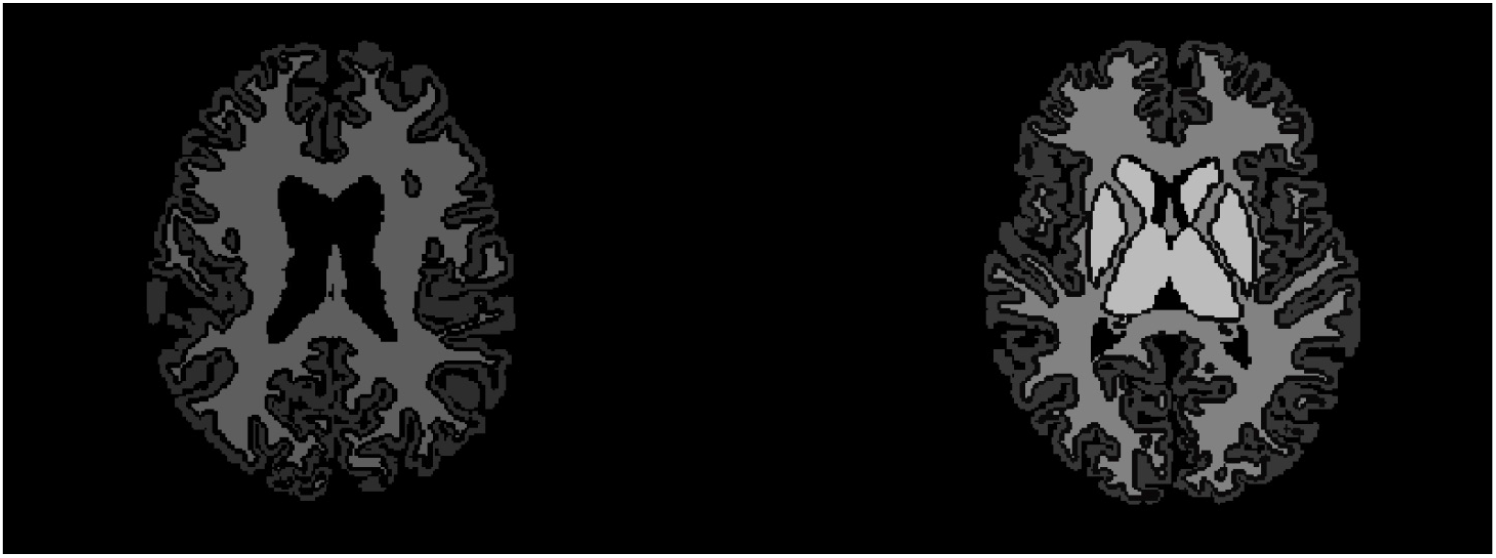
Whole-brain *K*_*i*_ medians per tissue provide scan-level summaries while preserving the GM>WM ordering.

### End-to-end transparency

Automated analysis is made auditable by exporting a compact visual summary of intermediate outputs (Figure 10) together with structured stage-level QC reports (described in Methods). These artifacts enable rapid identification of failed or atypical runs without manual per-frame inspection.

**Figure 10:**
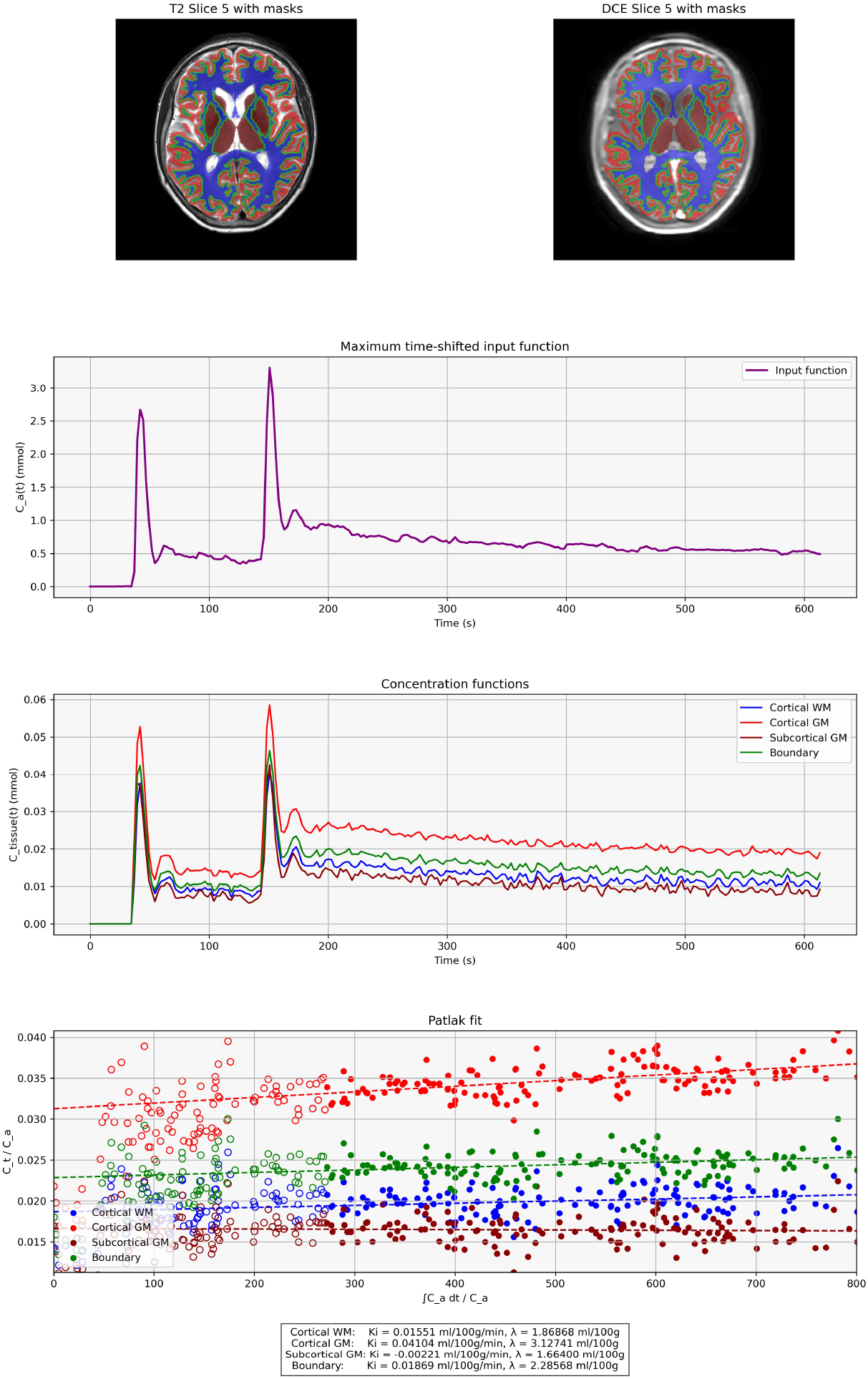
End-to-end transparency panel for a representative scan. We show segmentation, input functions, tissue curves, Patlak fits, and resulting *K*_*i*_ and *λ* (plasma volume, *v*_*p*_).

### Comparison to a reference implementation

We evaluated agreement between *p*-Brain and an established reference software workflow for Patlak-derived BBB permeability (*K*_*i*_). For this comparison, the reference workflow used manually defined arterial/venous input-function ROIs and tissue ROIs, and *p*-Brain was run in both manual and fully automatic modes.

Figure 11 compares gray- and white-matter Patlak-derived permeability (*K*_*i*_) summaries between *p*-Brain and the reference workflow across eight subjects with complete automatic outputs. In the reference workflow, tissue ROIs were manually placed in the right prefrontal cortex and right thalamus on a mid-brain slice (approximately slices 4–5 in the presented cohort), whereas *p*-Brain automatic measurements were derived from full-volume tissue masks (subcortical gray matter and cortical white matter). The between-method distributions showed no statistically significant differences in either tissue class.

**Figure 11:**
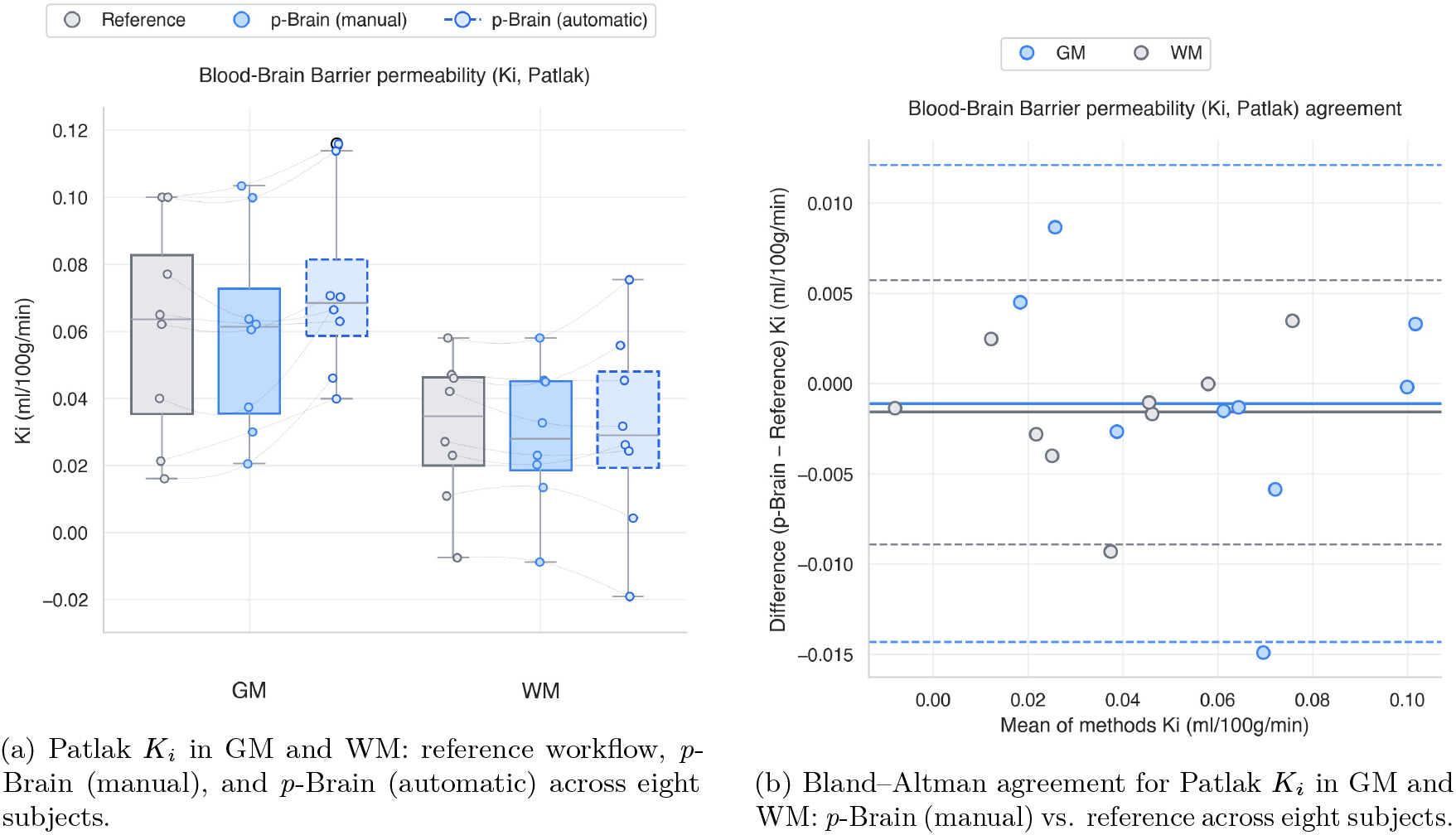
Automated comparison to an established manual reference workflow for whole-brain Patlak *K*_*i*_. Left: between-method distributions; right: Bland–Altman agreement.

Agreement was quantified using Pearson correlation and Bland–Altman analysis. For Patlak *K*_*i*_, agreement with the reference workflow was high (GM: *r* = 0.978, bias = − 5.11 *×* 10^−4^ mL*/*100g*/*min, 95% limits of agreement [−0.0141, 0.0131]; WM: *r* = 0.987, bias = −2.22 *×* 10^−3^ mL*/*100g*/*min, 95% limits [−0.00896, 0.00453]), indicating small systematic offsets and tight subject-level agreement for scan-level *K*_*i*_ summaries in this dataset.

In addition, voxelwise comparisons against the reference MATLAB program showed very good agreement for the perfusion map outputs and for the *T*_1_/*M*_0_ fitting outputs on a representative slice (Appendix Figures 20 and 21). In these comparisons, *p*-Brain was run in fully automatic mode, whereas the reference program relied on manual steps.

## 3 Discussion

The present work demonstrates that *p*-Brain can replace a traditionally manual, multi-software workflow for dynamic contrast-enhanced MRI with a reproducible automated pipeline. Direct agreement with reference implementations indicates that the automation does not sacrifice quantitative fidelity while eliminating operator-dependent variability that has historically hindered longitudinal and multi-site studies of blood–brain barrier physiology (Varatharaj et al., 2024a; Cramer et al., 2023a). By orchestrating segmentation, pharmacokinetic modeling, and multi-scale reporting in a single run, *p*-Brain offers a practical path toward routine deployment of quantitative DCE-MRI in research hospitals and, eventually, clinical radiology departments. Beyond providing reviewable figures, the pipeline exports analysis-ready outputs across scales—voxelwise NIfTI maps, parcel- and tissue-level summaries, cohort-level atlas projections, and scan-level whole-brain descriptors—together with structured logs and runtime metadata. This design supports downstream statistics and sharing across centres without requiring bespoke post-processing scripts.

The comparison to the reference workflow highlights expected sources of between-method variation, including differences in numerical implementation (e.g., optimization details) and the operator dependence of manually drawn ROIs. ROI definition and size can also influence stability: as ROI size increases, pharmacokinetic estimates generally become more stable due to averaging over noise and partial-volume effects, whereas *K*_*i*_ can appear inflated when estimated from small hand-drawn tissue ROIs compared with larger, atlas-derived tissue masks. Despite these factors, the observed bias and limits of agreement indicate close concordance of scan-level *K*_*i*_ summaries in this technically uniform dataset.

Our approach contrasts with previous semi-automated toolkits that still require labor-intensive arterial input selection, ROI drawing, or custom scripting to obtain voxel-wise and parcel-wise summaries. For example, ROCKETSHIP and related frameworks provide modular modeling utilities but expect considerable user supervision and do not guarantee harmonized outputs across centers (Barnes et al., 2015). Similarly, recent deep learning pipelines for tumor imaging often focus on specific pathologies and omit generalized whole-brain reporting or residue analysis (Nalepa et al., 2020). By comparison, *p*-Brain integrates CNN-based vascular and tissue segmentation, Patlak modeling, and model-free deconvolution to deliver BBB leakage, perfusion, and microvascular dispersion metrics in a unified exportable format.

In addition to the open-source Python implementation, we distribute a macOS desktop application that provides project organisation, batch job management, and responsive end-to-end execution on local machines, lowering the barrier to routine use by non-expert users while keeping sensitive data on-device.

The value of this integration is evident in the derived residue-based parameters. Capillary transit-time heterogeneity and mean transit time summarize complementary aspects of microvascular transport that influence oxygen delivery, and their inclusion enables interrogation of subtle hemodynamic alterations linked to vascular reserve and metabolic efficiency (Larsson et al., 2017b; Vestergaard et al., 2023b; Angleys et al., 2015). Because these summaries are produced alongside whole-brain and parcel medians, the same outputs can be reused for longitudinal monitoring, clinical decision support, or cross-cohort harmonization without revisiting the raw DCE data.

Automated anatomical segmentation underpins many of these advances. FastSurfer yields consistent gray- and white-matter parcellations within minutes (Henschel et al., 2020), and the released CNN models for vascular ROI extraction extend this principle to arterial input and venous outflow definition (Tireli, 2025). The resulting standardization reduces the learning curve for non-expert operators and mitigates variability arising from manual ROI placement (Cramer et al., 2023a). These gains are particularly relevant in large-scale studies of aging or neurodegeneration, where mild BBB leakage and perfusion changes must be detected against a backdrop of small effect sizes (Verheggen et al., 2020).

Finally, the pipeline preserves flexibility for future methodological advances. Because *p*-Brain exports intermediate concentration time series and segmentation masks, investigators can incorporate alternative kinetic models or machine learning predictors without reimplementing the ingestion and QC stages. Incorporating sequence harmonization and uncertainty quantification are natural next steps to improve robustness across sites and patient populations.

### Limitations

Despite these strengths, several practical constraints remain. First, the CNN models for anatomical and vascular segmentation were trained on relatively homogeneous data and can underperform in the presence of atypical anatomy, implants, or extensive pathology as well as sequences with varying contrast relationships, voxel dimensions or number of slices. When automated ROI extraction is not desired or is unsuitable for a given dataset, *p*-Brain can also be run with manual ROI delineation (vascular and/or tissue ROIs) via the GUI. For the segmentation used here, and as default, FastSurfer itself notes reduced reliability around large lesions, making visual inspection or targeted fine-tuning necessary when transferring to outlier populations (Henschel et al., 2020). Second, accurate kinetic modeling assumes physiologic regimes that may not hold universally. Patlak analysis presumes negligible backflux, and deconvolution-based perfusion estimates can be sensitive to input-function quality, bolus timing, and signal-to-noise; these assumptions can break in severe inflammatory states or other atypical physiology, biasing permeability and perfusion estimates (Varatharaj et al., 2019). Incorporating model selection logic, uncertainty quantification, or Bayesian approaches remains future work.

Motion and acquisition variability present additional challenges. Misregistration between the structural volume and the DCE time series directly perturbs the arterial input function and parcel propagation, degrading voxel-wise accuracy (Cramer et al., 2023a). Although the current pipeline performs rigid alignment and logs QC metrics, severe head motion or low signal-to-noise ratio datasets may still require reacquisition or more sophisticated correction. Finally, like most data-driven systems, *p*-Brain inherits domain shift sensitivity: scanner upgrades, protocol adjustments, or application to pediatric and geriatric cohorts can alter intensity distributions and vascular dynamics. Systematic validation across vendors and longitudinal recalibration of the CNN components and kinetic priors are essential before broad clinical deployment (Varatharaj et al., 2024a).

## 4 Materials and methods

### 4.1 Participants and MRI acquisition

We analysed DCE-MRI data acquired at Copenhagen University Hospital (Rigshospitalet) in a study of healthy participants. All participants provided written informed consent. Ethical approval was granted by the Danish National Research Ethics Committee (H-21047857) and all procedures complied with the Declaration of Helsinki.

All MRI data were acquired on a Philips 3T dSTREAM Achieva scanner with a 32-channel head coil. A high-resolution 3D *T*_1_-weighted turbo field echo sequence was acquired with TE = 5.13 ms, TR = 11.18 ms, flip angle = 8^◦^, and voxel size ≈ 0.67mm *×* 0.67mm *×* 0.70mm.

The dynamic contrast-enhanced acquisition was a 4D time series with 256 × 256 × 10 × 250 samples (ten slices, 250 time points), voxel size ≈ 0.898mm × 0.898mm × 9.50mm, echo time TE = 1.93 ms, and temporal sampling Δ*t* ≈ 2.463 s (total duration ≈ 616s). Although evaluated on this specific protocol, *p*-Brain does not assume a fixed DCE geometry: spatial dimensions and temporal sampling are read from NIfTI headers and JSON sidecars when available, with configurable overrides via environment settings to support other acquisition geometries. The pipeline supports both single- and double-bolus injections; in our data, the blood signal can exhibit two bolus peaks, and the default settings are tuned to this regime.

For baseline relaxation mapping, an inversion-recovery series was acquired at inversion times *T*_*I*_ ∈ {120 ms, 300 ms, 600 ms, 1000 ms with matching in-plane geometry to the DCE slices. A gadolinium-based contrast agent was administered intravenously according to the local clinical/research protocol.

### 4.2 Workflow overview

We summarize the end-to-end workflow in Figure 12. The gray shading marks fully-automatic steps. White boxes run with assisted automation and may, if desired, require human input for ROI drawing and review.

**Figure 12:**
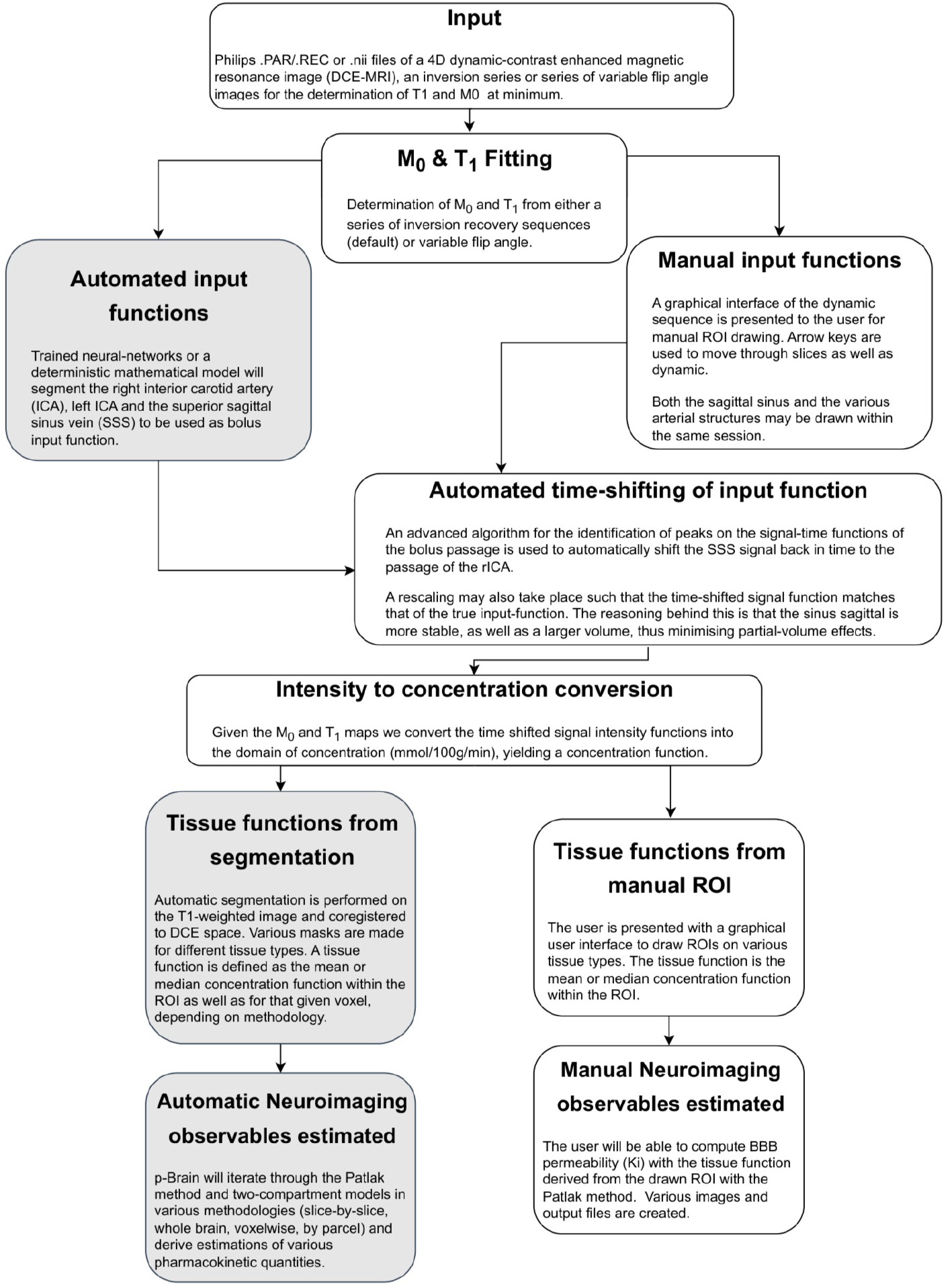
End-to-end *p*-Brain workflow. From *T*_1_*/M*_0_ fitting and automated AIF/VOF extraction to signal–to–concentration conversion, tissue-curve aggregation (manual or segmentation-based), and pharmacokinetic modeling (Patlak; optional deconvolution), yielding voxelwise, regional, slice-wise, and whole-brain outputs. The gray boxes denote fully automatic steps and white boxes indicate user-assisted stages.

The pipeline is organized into the following steps (see Figure 12):

1. **Inputs**. Required sequences are an inversion-recovery series for T_1_*/M*_0_ and a 4D DCE-MRI time series. A 3D *T*_1_-weighted sequence will allow the segmentation module to report outputs per parcel or tissue.
2. **Preprocessing**. Standard clinical imaging inputs (e.g., DICOM; NIfTI) are ingested and aligned to a common geometry. Structural volumes are rigidly aligned to the DCE acquisition. Temporal sampling remains as acquired. Further spatial corrections and intensity normalization are omitted in the present build.
3. *T*_1_ **and** *M*_0_ **maps**. *T*_1_ and *M*_0_ are estimated voxelwise by nonlinear least squares. The implementation supports both inversion–recovery and saturation–recovery signal models; acquisition-specific timing parameters are described in the *Participants + acquisition* section.
4. **Input functions**. The internal carotid artery (default reference: rICA) and the superior sagittal sinus are identified by the CNN-based slice classifier and ROI segmentation models when the fully automatic workflow is selected. Manual delineation of vascular ROIs is also supported via the GUI when AI-based ROI extraction is not used. By default, modelling uses an SSS-derived effective input function (TSCC) constructed by aligning the venous curve to the selected ICA reference, with an optional rescaling to account for transit delay and dispersion; the ICA-derived arterial input function can be used directly if required.
5. **Tissue ROIs**. Tissue ROIs may be hand drawn within the presented user-interface. In the fully automatic mode, the segmentation module, by default FastSurfer (Henschel et al., 2020), runs an anatomical segmentation on the *T*_1_-weighted image whereafter the segmentation module delineates cortical, subcortical and cerebellar gray matter, white matter, brainstem, and optional boundary masks defined as the boundary between subcortical grey matter and cortical white matter. An affine transformation aligns these masks to DCE space.
6. **Signal to concentration**. The pipeline converts signal–time curves into gadolinium concentration using the fitted *T*_1_ and *M*_0_ maps together with the acquisition timings from frames to seconds. Built-in safeguards reject invalid logarithmic terms.
7. **Modeling**. Patlak analysis yields *K*_*i*_ and *v*_*p*_, both available from manual analysis. Model-free Tikhonov deconvolution provides CBF, mean transit time (MTT), and capillary transit-time heterogeneity (CTH). The framework is designed to be extensible, enabling additional kinetic models to be incorporated as needed.
8. **Outputs**. The pipeline produces quantitative outputs at multiple scales, including voxelwise maps, regional metrics derived from anatomical parcellations, slice-wise summaries, and whole-brain summaries. Outputs are exported in standard research-friendly formats for direct use in statistical workflows. Parcellated summaries for CBF, MTT, and CTH are produced analogously to *K*_*i*_, using FastSurfer anatomical labels propagated into DCE space.
9. **Quality control and auditability**. Each stage emits reviewable artifacts (figures, masks, maps) and a structured quality control (QC) report that checks the presence and basic validity of its expected outputs (e.g., non-empty ROI masks, segmentation outputs, modelling summaries). QC is stage-scoped and can optionally be enforced to stop a run when a critical output is missing.

Each component of the workflow is detailed below in compact form, with full mathematical derivations and implementation settings provided in the Supplementary Methods.

### 4.3 *T*_1_ and *M*_0_ estimation

Voxelwise *T*_1_ and *M*_0_ were estimated from the inversion–recovery (IR) dataset by nonlinear least squares (Branch et al., 1999; Coleman and Li, 1996). For compatible protocols, *p*-Brain can alternatively estimate baseline *T*_1_*/M*_0_ from a variable flip-angle (VFA) spoiled GRE/SPGR series (multiple flip angles), using per-series repetition time and flip-angle values from the acquisition protocol; this option was not used for the data analysed for the presented data however.

For inversion times *T*_*I*_ = {*T*_*I*,1_, …, *T*_*I,N*_ } and observed voxel signals *y*_*i*_, the expected signal was expressed as

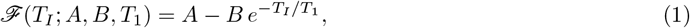

a form corresponding to the standard inversion–recovery signal model

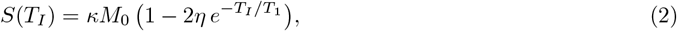

where *η* is the inversion efficiency (typically ≤ 1) and *κ* is a receive–gain scale. In *p*-Brain we fit Eq. 1 and do not estimate *η* or *κ* separately; their effects are absorbed into *A* and *B* (with *A* = *κM*_0_ and *B* = 2*ηκM*_0_). This model describes recovery from inverted magnetisation (− *M*_0_) to equilibrium. Inversion times for the dataset analysed here are given in *Participants and MRI acquisition*.

Parameter estimation proceeded voxelwise by minimising the residual energy

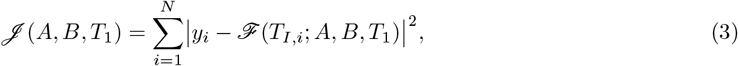

subject to *T*_1_ ∈ [100, 6000] ms. In the released implementation, *A* and *B* are constrained to be positive; the signed IR signal can cross zero around the null point (*T*_*I*_ ≈ *T*_1_ ln 2) when *B > A*. This ensures unbiased *T*_1_ estimates across tissues where the acquisition samples both sides of the inversion null. Initial guesses were *A* = max_*i*_ *y*_*i*_, *B* = *A/*2, and *T*_1_ = 750 ms; voxels with negligible signal were excluded.

The fitted maps 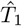 and Â retain the geometry of the original inversion–recovery images (Figs. 13 and 14). An *M*_0_ proxy can be recovered as 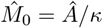 when receive calibration is available. These maps provide baseline relaxation parameters required for quantitative DCE-MRI analysis, enabling conversion of signal intensity to gadolinium concentration.

**Figure 13:**
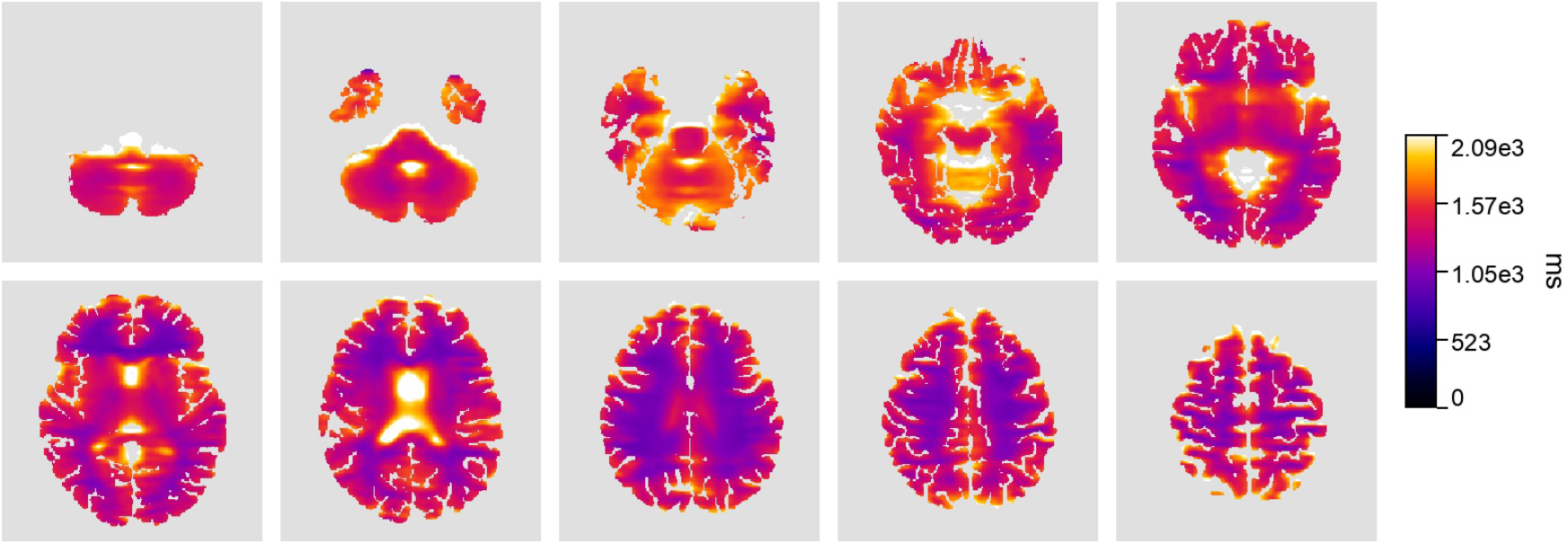
Voxelwise *T*_1_ (ms) values from a representative ten-slice inversion-recovery dataset.

**Figure 14:**
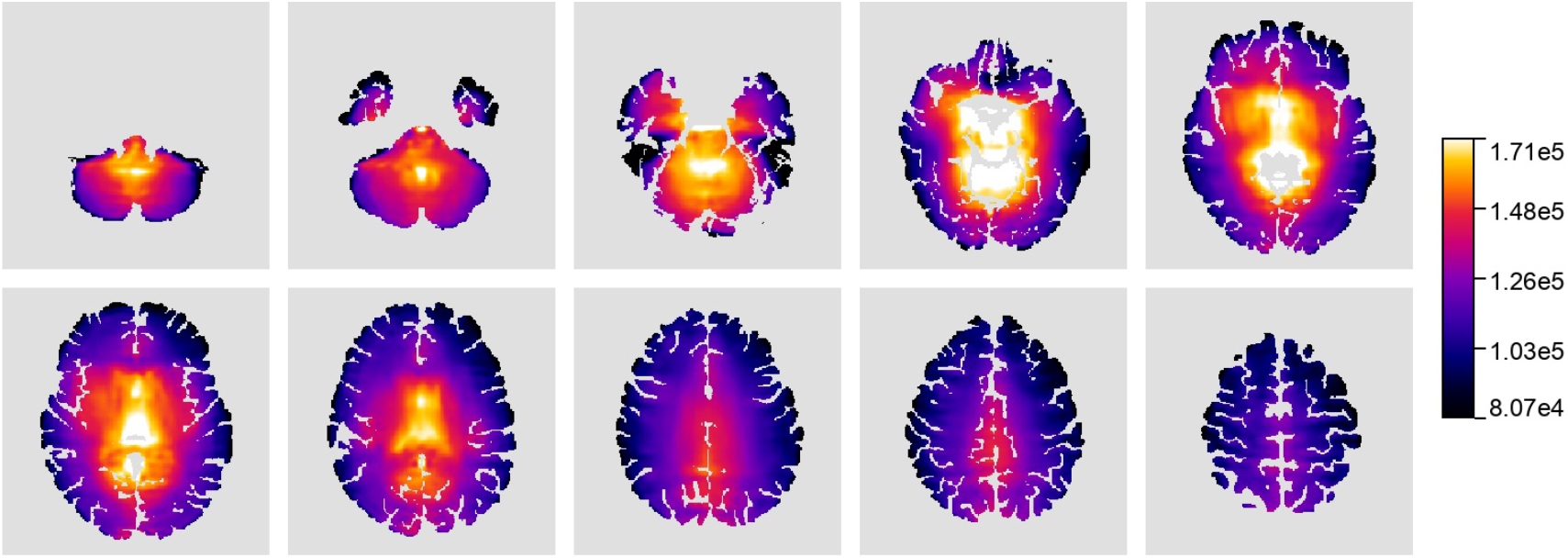
Voxelwise *M*_0_ magnetisation values from a representative ten-slice inversion-recovery dataset.

**Figure 15:**
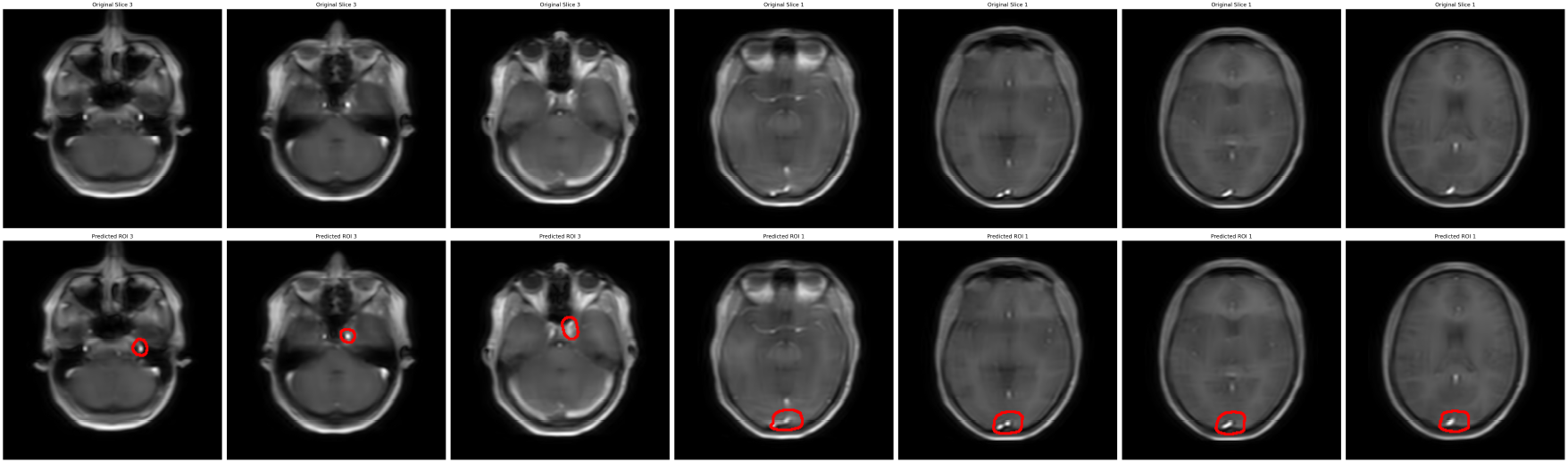
CNN slice selection and ROI delineation for the right internal carotid artery and superior sagittal sinus.

**Figure 16:**
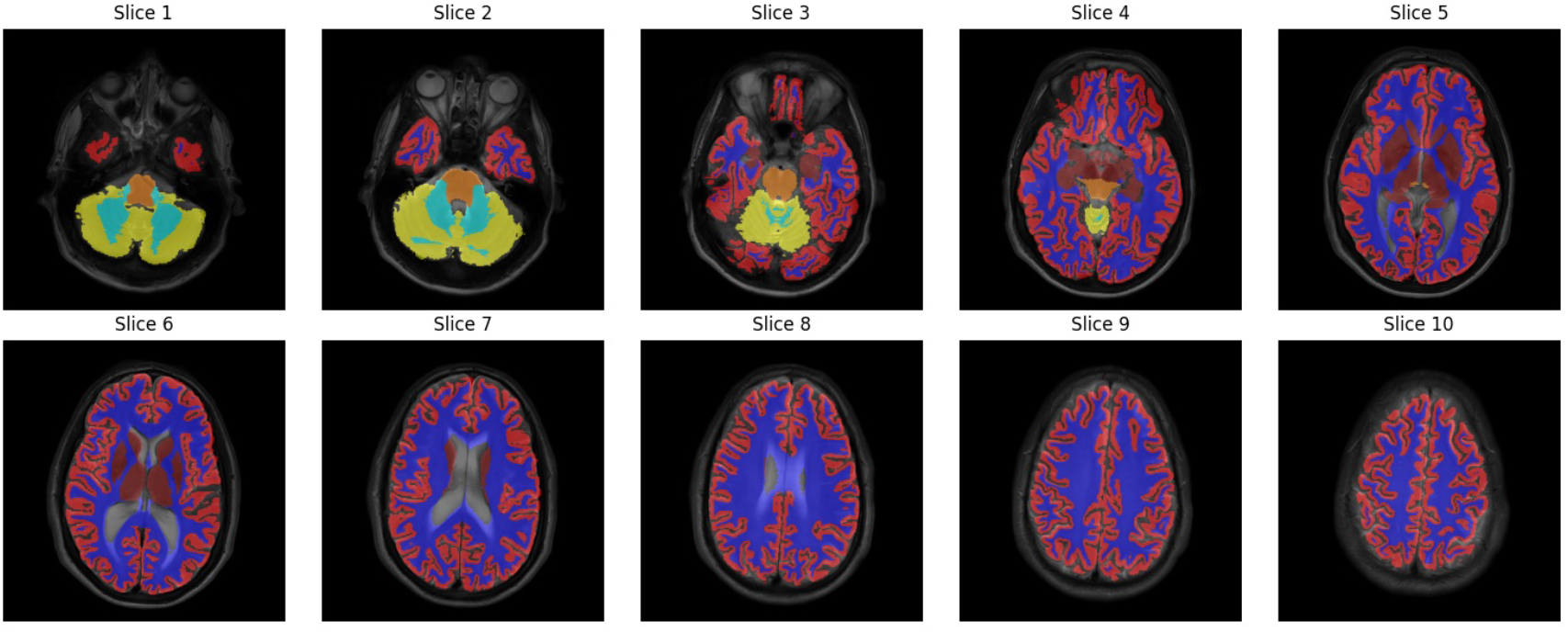
Anatomical segmentation overlaid on an axial T1-weighted image. Cortical gray matter (red), subcortical gray matter (dark red), white matter (blue), brainstem (orange), cerebellar white matter (light blue), and cerebellar gray matter (yellow).

In the current released implementation, voxelwise gadolinium concentration curves are derived using the fitted *T*_1_ and *M*_0_ maps via the saturation-recovery-based conversion (see *Signal-to-concentration conversion*).

### 4.4 Vascular and tissue ROI extraction

Regions of interest (ROIs) support two core tasks in the pipeline: (i) deriving vascular input functions (arterial and venous) and (ii) aggregating tissue concentration–time curves for downstream modeling.

For the arterial input function (AIF), we extract the signal from an internal carotid artery (default: rICA; lICA can be selected if required, e.g., if the rICA curve is degraded), which provides a robust measure of contrast arrival to the brain. The superior sagittal sinus (SSS) serves as the venous output function (VOF), reflecting contrast washout. As described in Workflow Step 4, the venous curve is aligned to the arterial peak via cross-correlation and amplitude rescaling to compensate for transit delays and dispersion (Cramer et al., 2023b; Manning et al., 2021; Thrippleton et al., 2019). In the fully automated implementation, we employ a two-stage CNN-based approach:

1. **Slice Classification CNN:** Identifies MRI slices containing relevant vascular and tissue structures based on intensity profiles and anatomical context.
2. **ROI Segmentation CNN:** Precisely delineates ROIs around the ICA and SSS for reliable extraction of input functions (for lICA selection, the ICA model can be applied on left–right mirrored slices and the predicted mask mirrored back).

Training used manually labelled examples of the ICA and SSS. The ROI segmentation networks are 2D U-Nets (Ronneberger et al., 2015); training and evaluation details are provided in the Supplementary Methods. Trained model weights are distributed with the software and are also available on Zenodo (Tireli, 2025).

Manual ROI definition is supported via the GUI (hand-drawn ROIs with user-selected slices) when AI-based ROI extraction is not used or when additional review is required.

For automated tissue ROI extraction, we employ anatomical segmentation using FastSurfer (Henschel et al., 2020) and FSL (Jenkinson et al., 2012), which generate masks for anatomical parcels directly from segmentation and further *p*-Brain aggregates cortical and subcortical gray matter, white matter, cerebellum, and brainstem through fully automated processing.

A boundary mask between gray and white matter is computed by overlapping dilated masks and eroding the intersection to a few-voxel thickness^2^. All segmentation labels are then propagated into DCE-MRI space using affine registration, ensuring reliable alignment despite patient motion or anatomical variability. These propagated masks provide the foundation for accurate voxelwise, regional, and parcellated pharmacokinetic analyses.

### 4.5 Signal–to–concentration conversion

DCE-MRI yields a four-dimensional dataset in which each voxel has a signal–time curve *S*(*t*). After defining arterial, venous, and tissue ROIs, we extract representative curves for modeling. For vascular ROIs, the default is to select the single brightest voxel within the mask (maximum over time) to reduce partial-volume contamination; alternatively, users can average the signal across ROI voxels. When using the max-voxel strategy, *p*-Brain also supports an *adaptive-max* variant in which the brightest voxel is re-selected independently at each time frame within the ROI to reduce sensitivity to in-plane motion. For tissue ROIs, we average voxelwise signals within the mask to reduce noise and improve stability.

We convert signal–time curves *S*(*t*) to gadolinium concentration 𝒞 (*t*) using the voxelwise *T*_1_ and *M*_0_ maps via the saturation-recovery conversion used in the released *p*-Brain implementation:

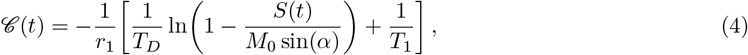

with *T*_1_ and *T*_*D*_ expressed in seconds, *α* denoting the *α*-pulse flip angle, prepulse-to-readout delay *T*_*D*_ = 120 ms (here for saturation recovery), and relaxivity *r*_1_ = 4 s^−1^ mM^−1^ (blood at 3 T). Sequence timing and flip-angle parameters were taken from the acquisition protocol. For the DCE protocol analysed here, the nominal readout flip angle was 30^◦^. To ensure numerical robustness, we clamp the logarithm argument to (0, 1).

For spoiled gradient-echo (SPGR) acquisitions with variable flip angles, *p*-Brain also supports a VFA-based conversion that first estimates *T*_1_(*t*) from the SPGR steady-state equation

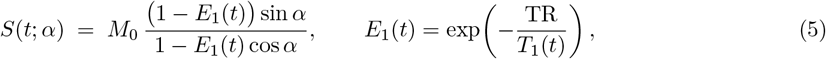

using two or more flip angles. A common linearisation writes

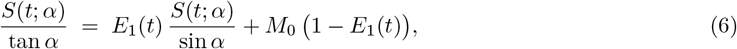

so that *E*_1_(*t*) (slope) and *M*_0_(1 − *E*_1_(*t*)) (intercept) can be estimated by regression across flip angles at each time point, yielding *T*_1_(*t*) = − TR*/* ln *E*_1_(*t*). Concentration then follows from the relaxivity relation 1*/T*_1_(*t*) = 1*/T*_1,0_ + *r*_1_ 𝒞(*t*), i.e.

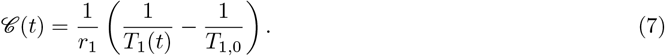

This VFA/SPGR branch is provided for compatible protocols but was not used for the data analysed here.

#### 4.5.1 Curve normalization and input-function selection

##### Baseline enforcement and curve scaling

After signal–to–concentration conversion, the pre-contrast concentration should be zero: gadolinium is absent before injection and any baseline fluctuations primarily reflect noise and modelling/fit error. *p*-Brain therefore applies a lightweight baseline correction to concentration–time curves (arterial, venous, and tissue) by subtracting a pre-bolus baseline estimate and enforcing a zero baseline window.

Let *c*_*i*_ denote the sampled curve at time index *i* and let *B* be the number of baseline frames (default *B* = 5). We estimate the pre-bolus baseline as the mean over the initial frames,

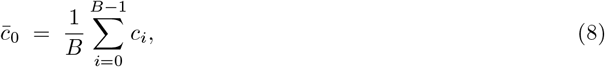

then baseline-correct the curve and enforce a zero baseline window,

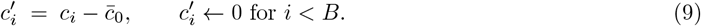

To reduce sensitivity to outliers and inter-voxel amplitude differences, we rescale by a robust deviation scale (95th percentile),

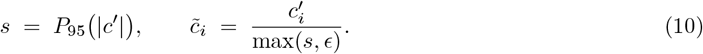

To optionally emphasise curve *shape* (timing and relative bolus features) over absolute amplitude in visualisations, we rescale by a robust percentile as above, yielding the behaviour illustrated in Figure 17. This rescaling can be disabled via configuration.

**Figure 17:**
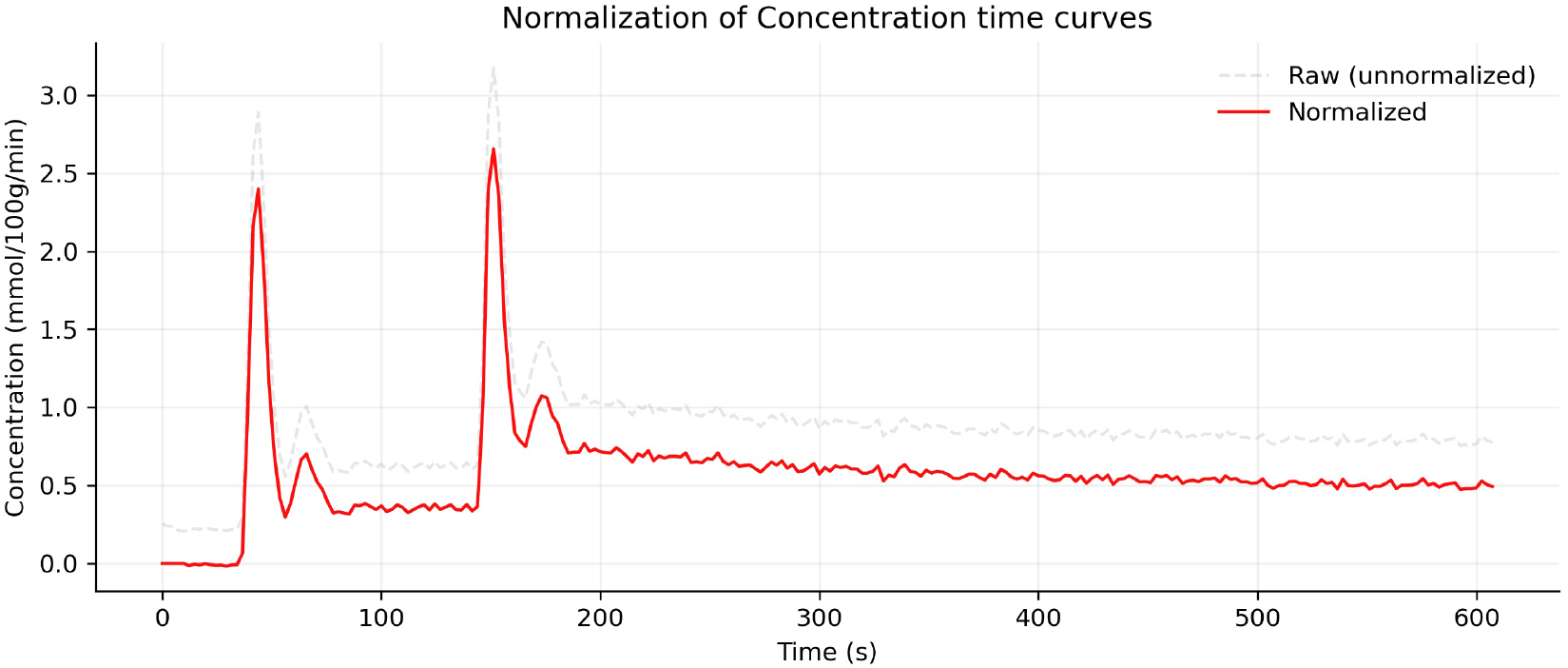
Example tissue concentration–time curve before and after the optional baseline/robust normalization used for diagnostics and visualization. Baseline frames are set to zero after subtracting the baseline mean, and the remaining curve is rescaled by a robust percentile to emphasise shape.

**Figure 18:**
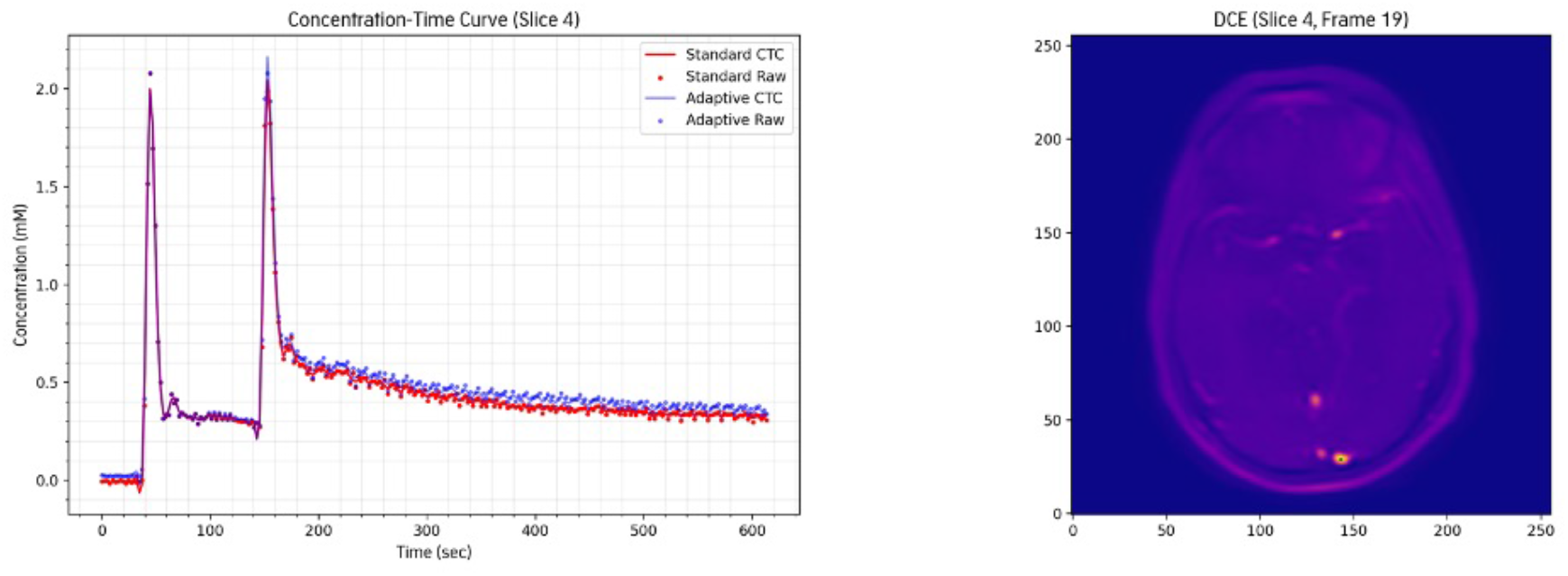
Time-shifted venous concentration–time curve for a double-bolus acquisition. The AIF is derived with adaptive signal tracking to correct minor in-plane motion. The concentration curves in this plot are passed through a Butterworth filter (Butterworth, 1930) for better interpretability.

**Figure 19:**
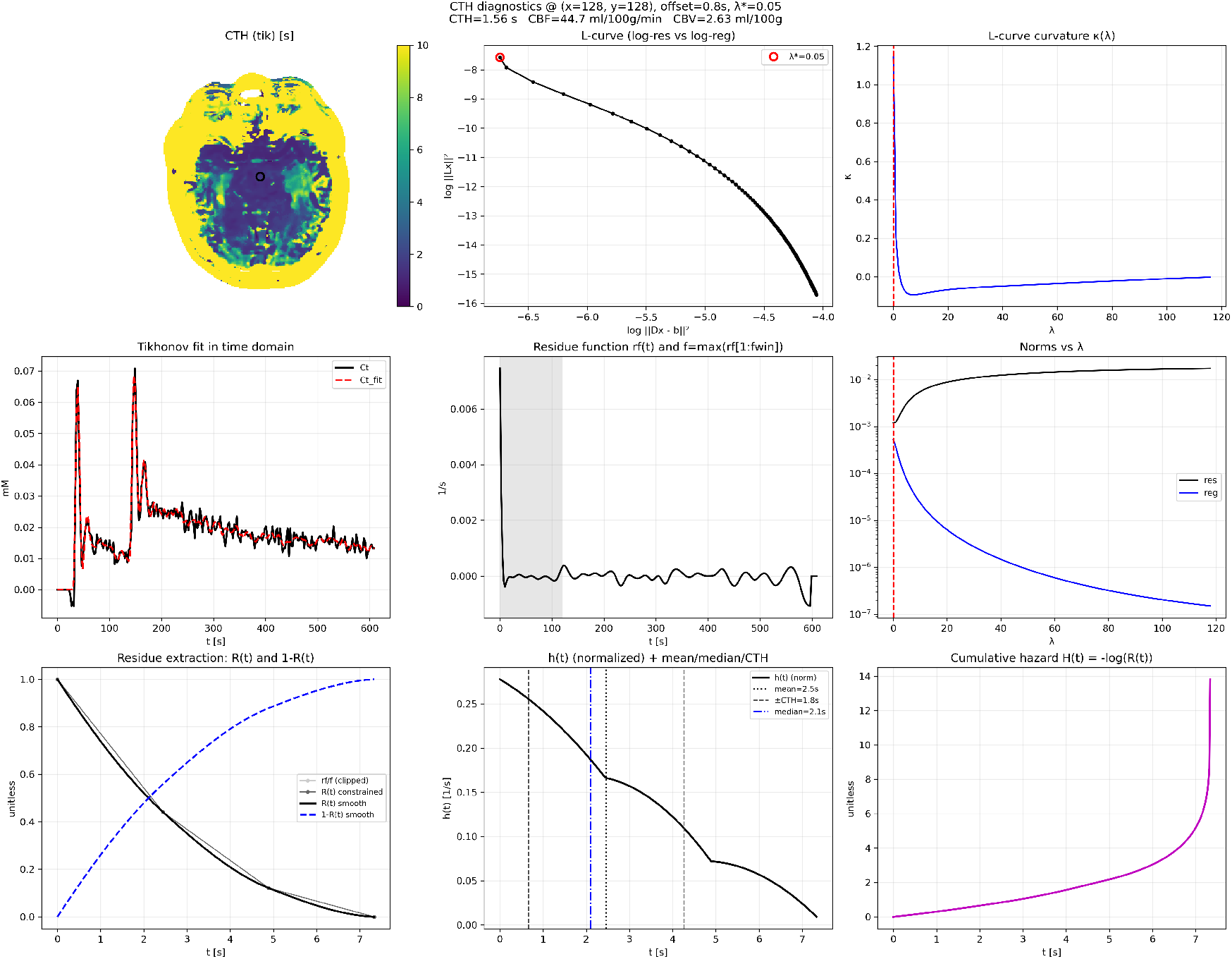
Example deconvolution diagnostics from the released *p*-Brain Tikhonov implementation for a representative voxel. The panel shows the L-curve (log residual norm vs log regularisation norm), curvaturebased selection of the optimal *λ*, the resulting time-domain fit, the reconstructed impulse response and constrained residue, the derived outflow distribution *h*(*t*), and the corresponding MTT/CTH summaries.

**Figure 20:**
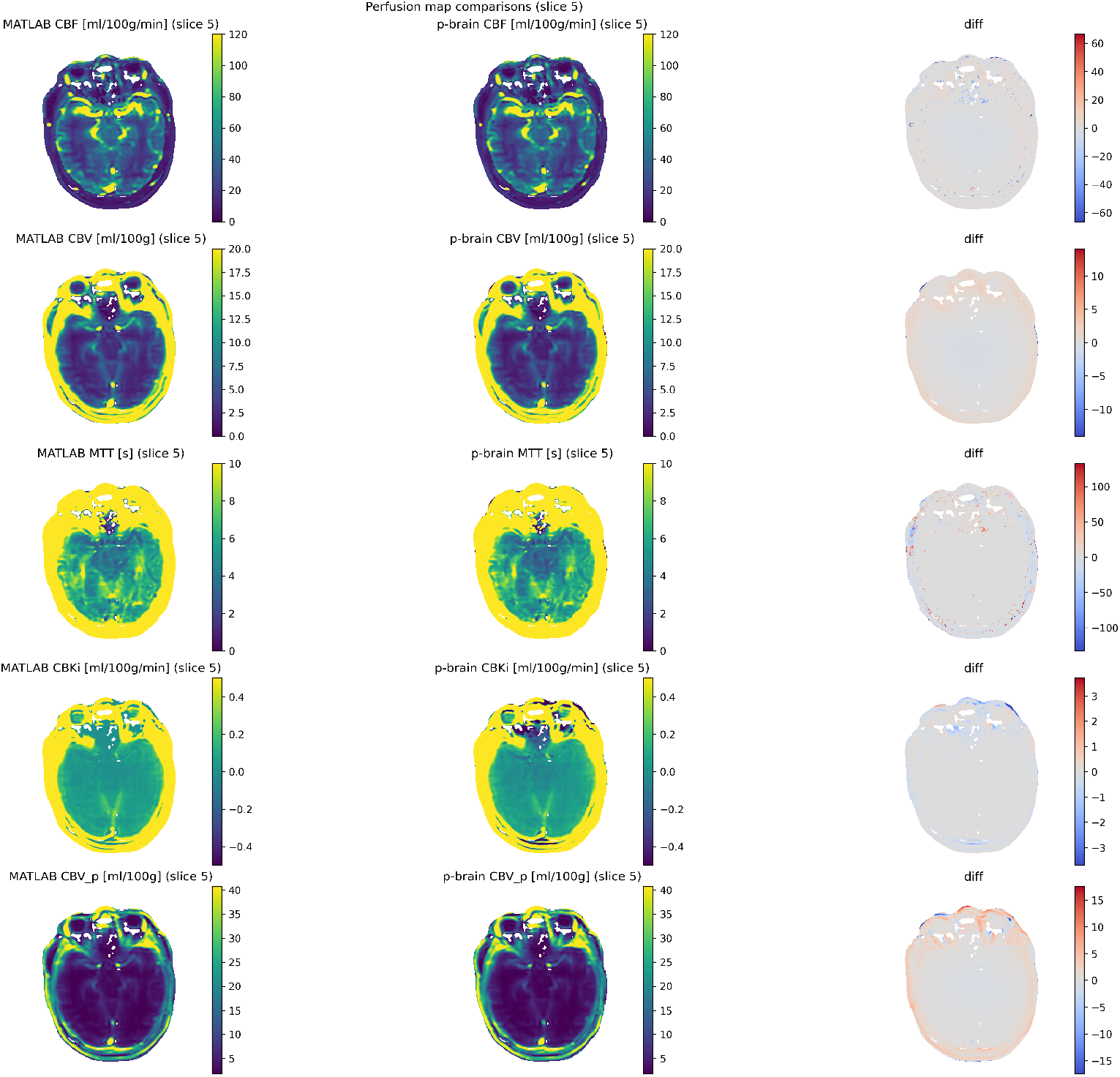
Voxelwise comparison of perfusion map outputs for a representative slice (slice 5): *p*-Brain (fully automatic) versus the reference MATLAB program (manual).

**Figure 21:**
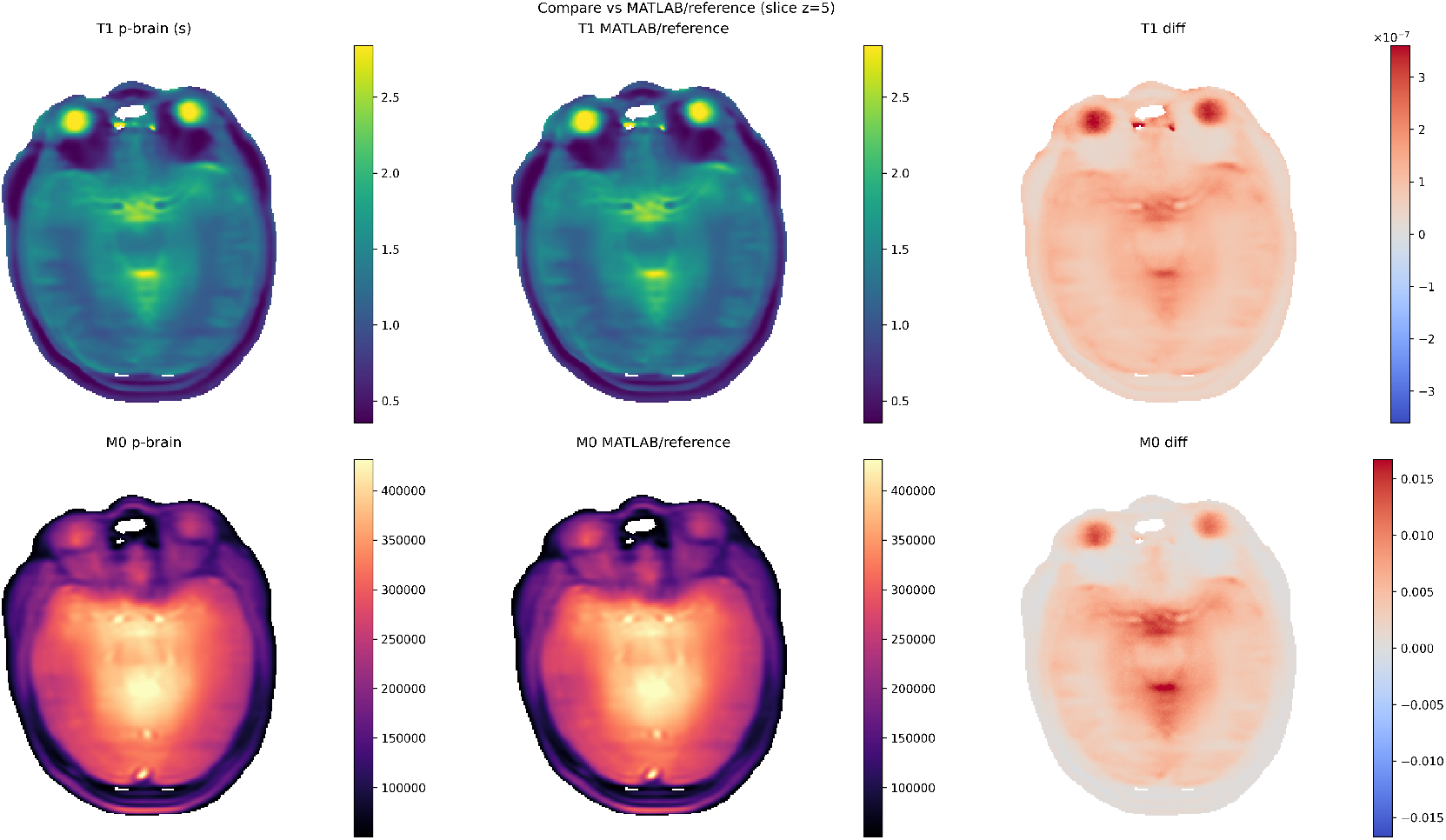
Voxelwise comparison of *T*_1_ and *M*_0_ fitting outputs for a representative slice (slice 5): *p*-Brain (fully automatic) versus the reference MATLAB program (manual).

**Figure 22:**
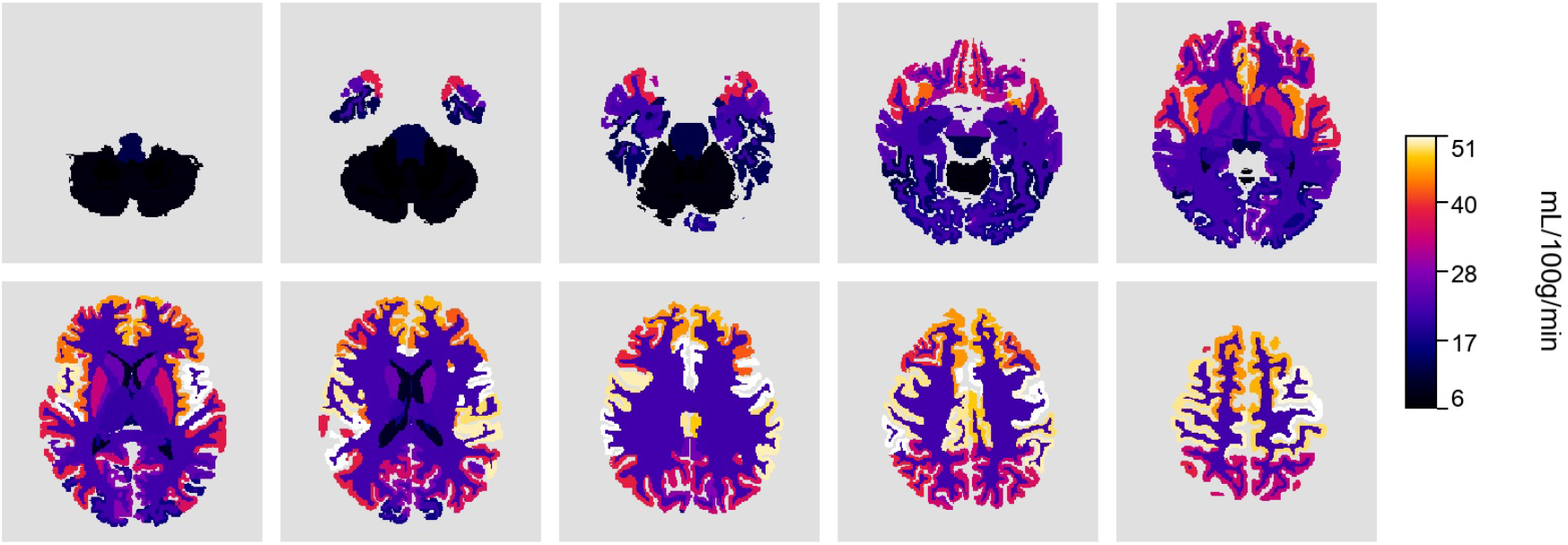
Cohort-mean CBF from model-free deconvolution. Units: mL*/*100g*/*min.

**Figure 23:**
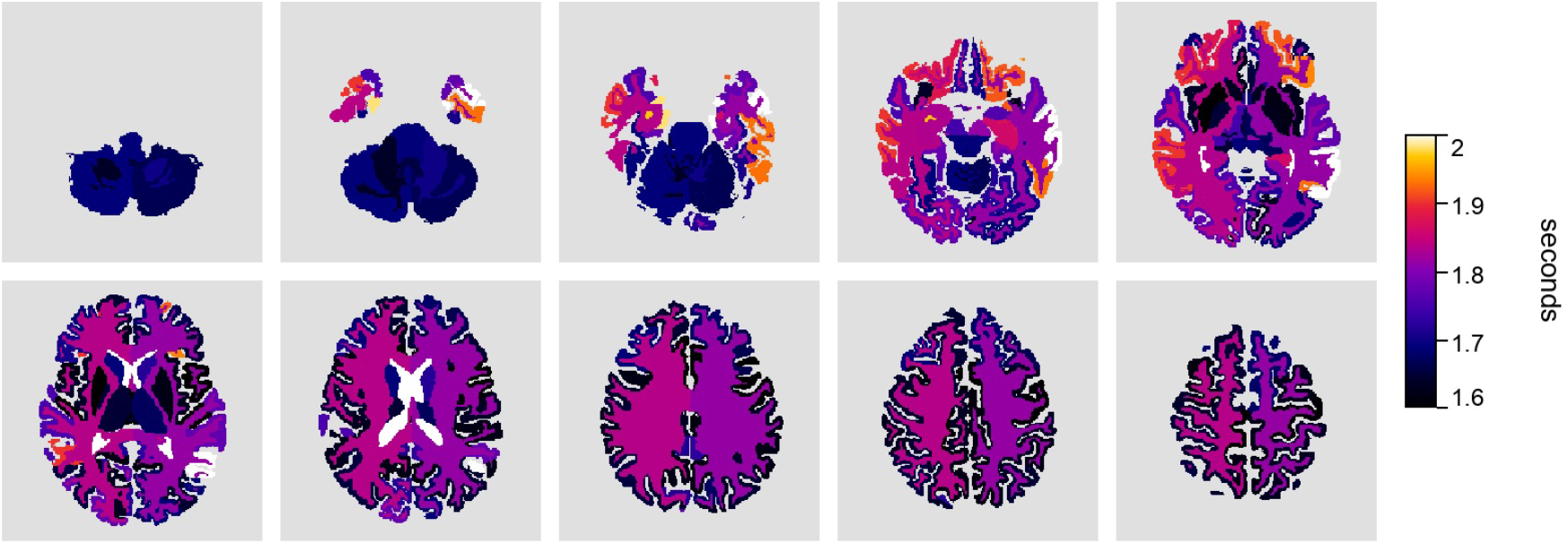
Cohort-mean capillary transit-time heterogeneity (CTH). Units: s.

**Figure 24:**
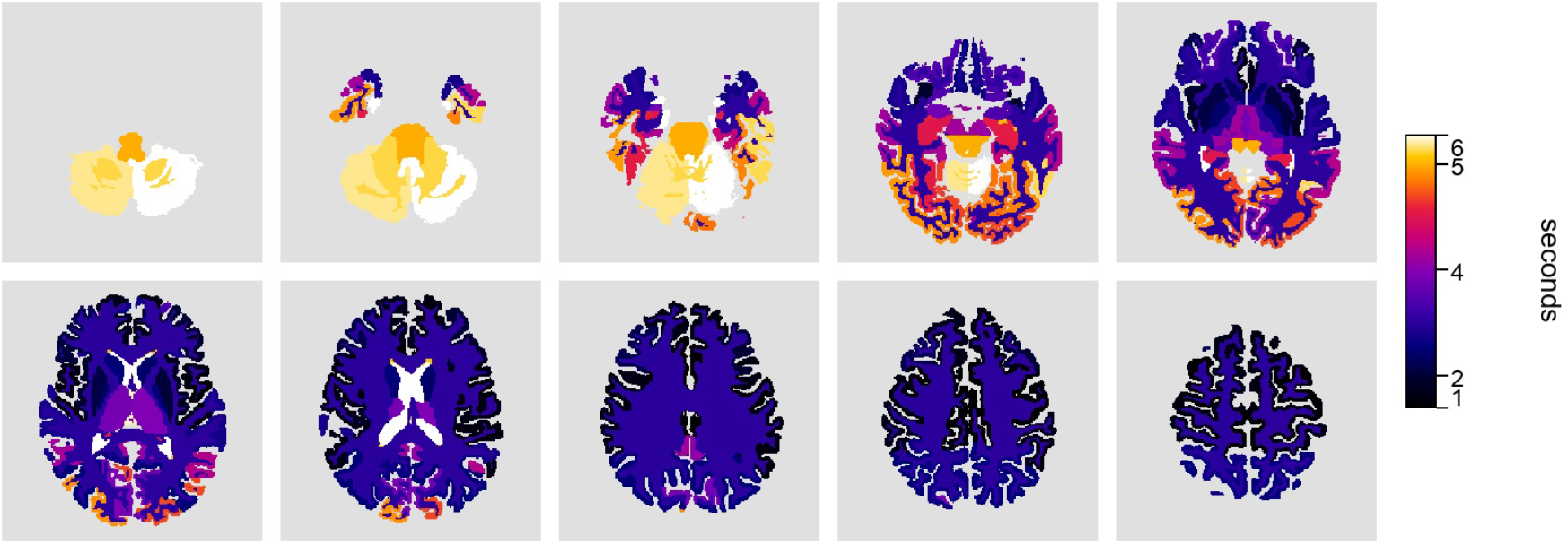
Cohort-mean mean transit time (MTT). Units: s.

**Figure 25:**
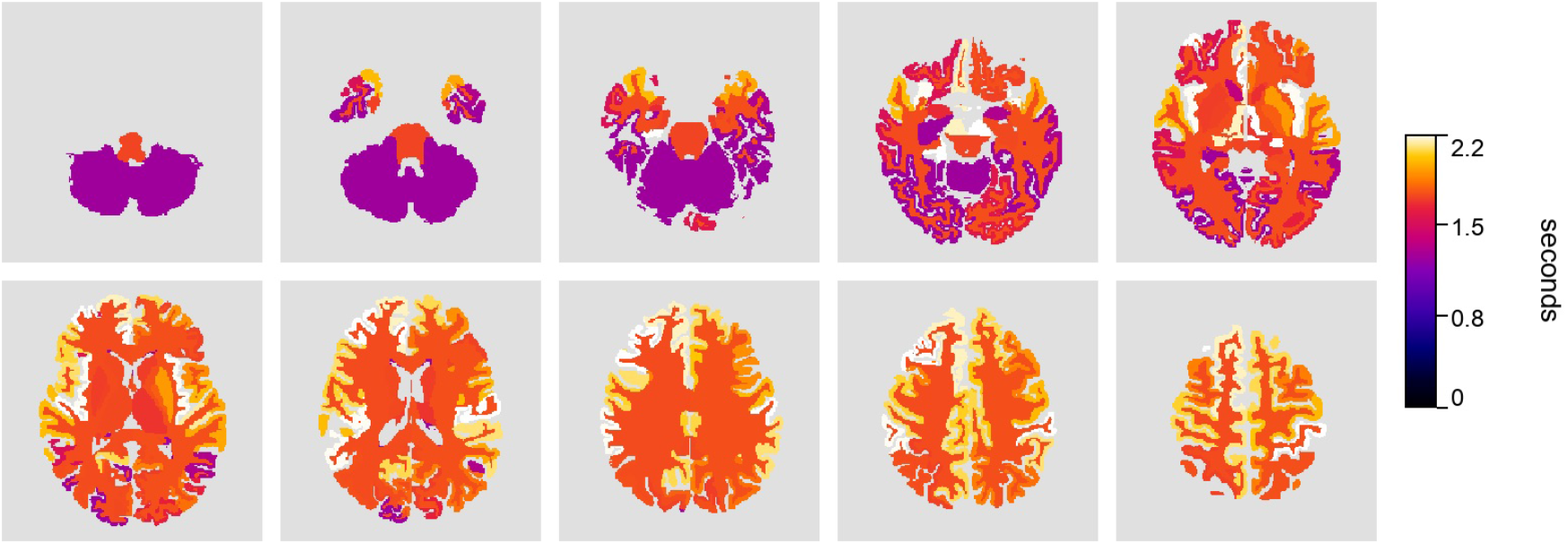
Parcel-level CTH in the representative participant. Units: s.

**Figure 26:**
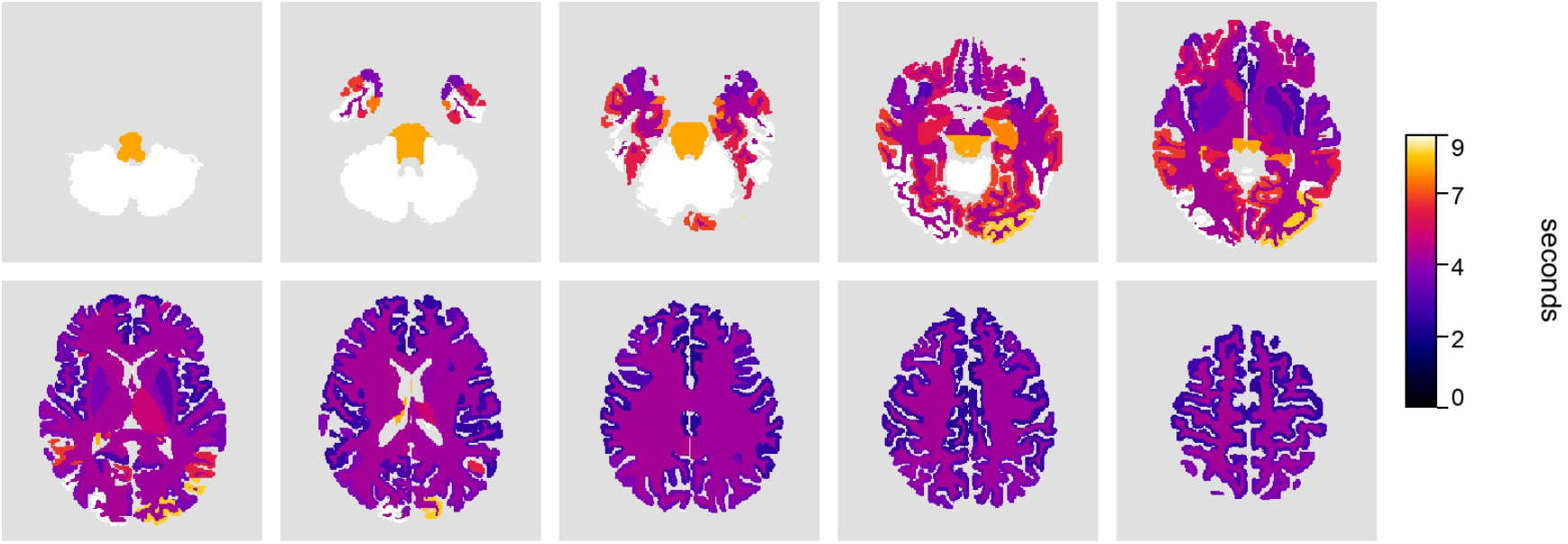
Parcel-level MTT in the representative participant. Units: s.

**Figure 27:**
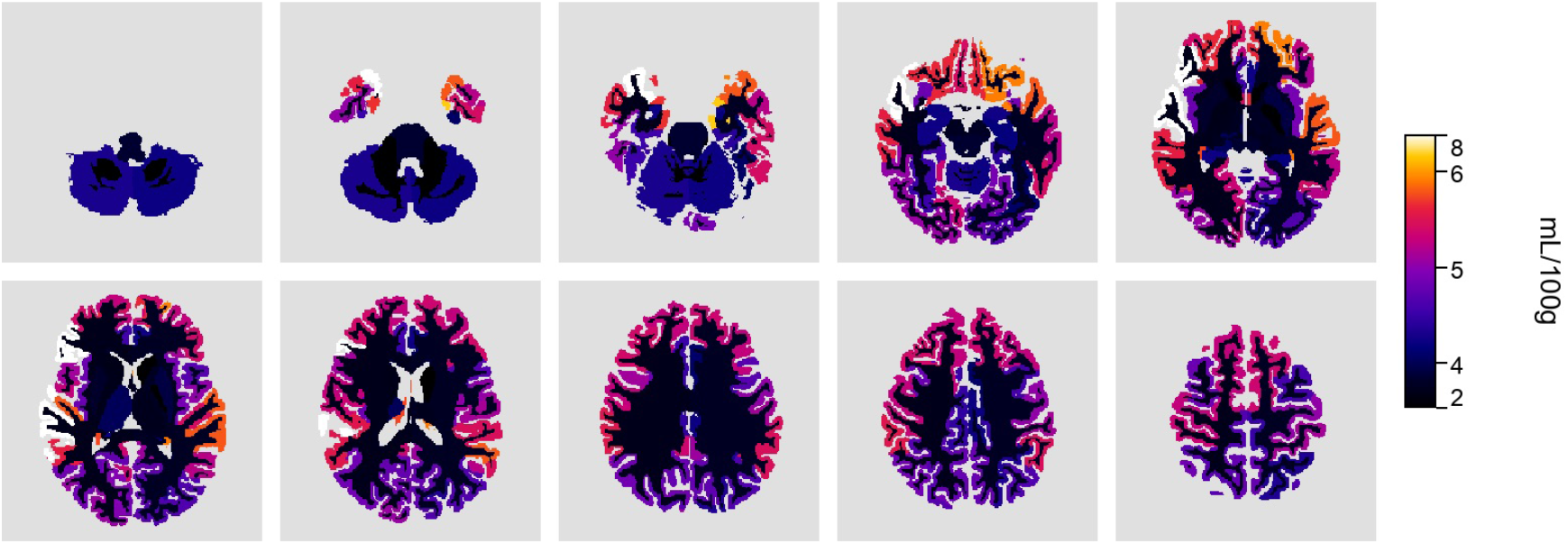
Parcel-level *v*_*p*_ in the representative participant.

##### Input-function selection (modelling)

For kinetic modelling, *p*-Brain uses an SSS-derived effective input function constructed by time-shifting the venous curve to a selected ICA reference (default: rICA; lICA can be selected if required, e.g., if the rICA curve is degraded) via cross-correlation peak alignment. An optional amplitude rescaling then produces a time-shifted (and optionally rescaled) concentration curve (TSCC). Rescaling can be disabled (time-shift only), or computed either by matching peak height (peak-based) or by matching the total curve “volume” (area-under-curve based). The ICA-derived arterial input function can also be used directly.

Input functions follow the automated extraction described in Workflow Step 4: the arterial curve is drawn from the right internal carotid artery and the venous curve from the superior sagittal sinus, with cross-correlation alignment and amplitude rescaling to compensate for transit delays and dispersion (Varatharaj et al., 2024b). We further apply adaptive signal tracking to mitigate in-plane motion when deriving the AIF.

### 4.6 Tracer–kinetic models and parameter estimation

#### 4.6.1 Patlak graphical analysis

We quantify low-level BBB leakage with the Patlak model, which assumes negligible backflux from the extravascular–extracellular space (EES) to plasma (*k*_2_ ≈ 0) and yields a linear relationship between tissue and input concentrations (Patlak et al., 1983). Let *C*_*a*_(*t*) denote the selected input function (default: TSCC; optional: ICA-derived AIF) and *C*_*t*_(*t*) the tissue concentration. The model reads

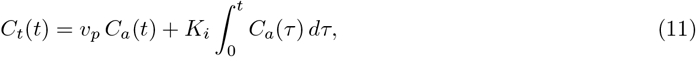

with *K*_*i*_ the influx constant and *v*_*p*_ the fractional plasma volume. Dividing by *C*_*a*_(*t*) gives a linear form,

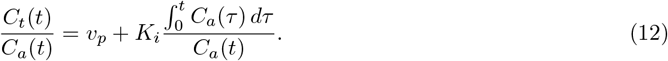

For discrete sampling at *t*_*i*_, define

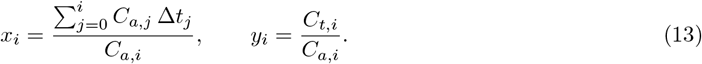

We fit *y*_*i*_ = *v*_*p*_ + *K*_*i*_*x*_*i*_ by ordinary least squares over the late (approximately linear) epoch of the Patlak plot; in the released implementation this is selected as the upper fraction of *x*_*i*_ (default: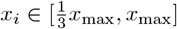) after excluding invalid samples (e.g., *C*_*a*_(*t*_*i*_) = 0 or NaNs), yielding

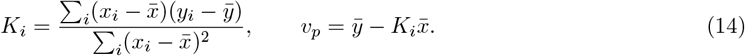

We report uncertainty as

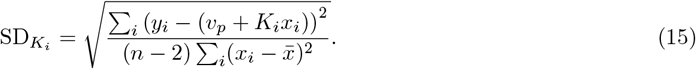

In the released *p*-Brain implementation, the cumulative integral in *x*_*i*_ is evaluated with a left Riemann sum on the acquired grid (with *x*_0_ = 0), i.e.

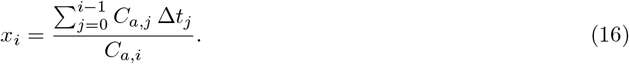

The fitted slope and intercept follow legacy reporting conventions: *K*_*i*_ is converted from s^−1^ to mL*/*100g*/*min by multiplying by 6000, and *v*_*p*_ is converted from a unitless fraction to mL*/*100g by multiplying by 100.

#### 4.6.2 Model-free deconvolution and microvascular transit metrics

We model the tissue concentration–time curve *C*_*t*_(*t*) as a convolution of the arterial input function *C*_*a*_(*t*) with a causal impulse response *g*(*t*) = *F R*(*t*), where *F* is cerebral blood flow and *R*(*t*) is the residue function with *R*(0) = 1:

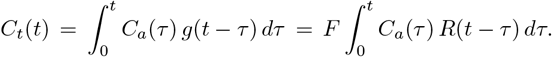

Sampling on a uniform grid *t*_*i*_ with step Δ*t*, recovering *g* from discrete measurements is ill-posed. In the released *p*-Brain implementation, we represent the impulse response in a cubic B-spline basis and solve for basis coefficients with Tikhonov regularisation. Let *B* be the B-spline basis matrix evaluated on the acquired grid, so that **g** ≈ *B*^⊤^**x***/*Δ*t*. With *A* the lower–triangular Toeplitz convolution matrix built from the (optionally time-shifted) input function, define *D* = *AB*^⊤^ and solve

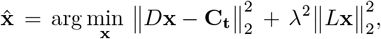

where *L* is a first-difference operator on the coefficient vector and *λ >* 0 the regularisation weight. Candidate *λ* values are evaluated on a fixed grid (default: 121 values spanning [0.05, *σ*_max_(*A*_0_)], where *A*_0_ is the Toeplitz matrix built from the unshifted input and *σ*_max_ its largest singular value). For each voxel, *λ* is selected by maximising the curvature of the L-curve in log 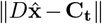 versus log 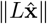 space. Optional per-voxel temporal offsets are handled by shifting *C*_*a*_(*t*) within the forward operator via a shape-preserving cubic interpolation; offsets can be quantised into small groups for efficiency.

##### Flow, blood volume, and transit metrics

After estimating 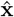, we reconstruct the impulse response **ĝ** and estimate flow as the early peak of the impulse response,

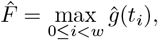

with a fixed peak window *w* (default: 50 samples). Cerebral blood flow is then reported in mL*/*100g*/*min using a density scaling (default tissue density *ρ* = 1.04 g mL^−1^); when the input function is plasma-derived, an additional (1 − Hct) factor is applied.

The released implementation reports an apparent distribution volume

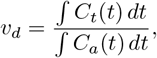

where the denominator uses the time-shifted input when offsets are enabled (numerical integration by the trapezoidal rule). Mean transit time is then computed as

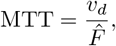

which is consistent with 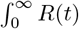 *dt* under the convolution model. The central volume relationship 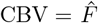 MTT therefore holds by construction.

##### CTH from the constrained residue

For microvascular dispersion, we form a normalised residue 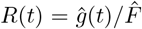 and apply stabilising shape constraints prior to differentiation: the residue is peak-aligned (so the impulse peak is at the start), renormalised to satisfy *R*(0) = 1, clipped to [0, 1], enforced to be non-increasing, and forced to taper smoothly to zero at the tail. We then compute

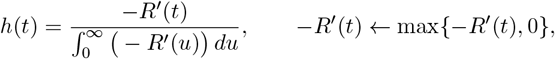

using finite differences on the acquired grid and trapezoidal-rule normalisation on midpoints. Finally,

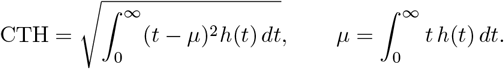

### 4.7 Quality control and end-to-end auditability

*p* rain is designed to produce standardized outputs while remaining auditable in batch operation. In addition to visual summaries (e.g., per-slice montage panels of masks, curves, and fits), the pipeline runs a lightweight quality-control (QC) layer after key stages. QC emits one structured report per stage with an overall status (pass, warning, or fail), together with brief messages that point to the relevant outputs for review. Optionally, QC can be enforced to stop a run when a stage reports failure.

The current QC checks are intentionally conservative: they verify that each stage produced its expected outputs and that key masks/curves are non-empty, rather than attempting to judge physiological correctness. Specifically, the following stage-level checks are performed:

- *T*_1_*/M*_0_ **fitting**. Checks that baseline relaxation mapping completed successfully and produced the expected parameter maps. Missing outputs yield fail.
- **Input functions**. Checks that arterial/venous ROIs were produced and are non-empty (i.e., contain voxels) so that AIF/VOF curves can be extracted. Unreadable outputs yield warning; missing or universally empty ROIs yield fail.
- **Time shifting**. Checks that the time-shift/alignment step produced readable selection metadata
- for the chosen ICA reference and (when configured) the TSCC alignment used for modelling. When modelling uses the arterial curve directly, missing TSCC metadata is downgraded to warning.
- **Segmentation**. Checks that the anatomical segmentation completed and produced the expected tissue/parcellation outputs for downstream aggregation. Missing segmentation yields fail.
- **Tissue curve export**. Checks that tissue concentration–time curves were exported for the configured tissue ROIs. Missing outputs yield fail.
- **Modelling**. Checks that model summaries were written and that the requested modelling outputs (summaries and, when enabled, maps) were produced. Missing summaries yield fail; missing optional maps yield warning.

In practice, QC reports complement the visual end-to-end montage (Figure 10) by highlighting runs that warrant closer review.

## 5 Conclusion

Our fully automated platform provides a standardized, reproducible framework for generating quantitative metrics of cerebral perfusion and permeability. Here we validate technical agreement of Patlak-derived *K*_*i*_ estimates in a uniform dataset; generalization across scanners, sites, pathologies, and clinical workflows remains to be established.

## List of Acronyms

AIF: arterial input function
BBB: blood-brain barrier
CBF: cerebral blood flow
CNN: convolutional neural network
CTH: capillary transit-time heterogeneity
DCE: dynamic contrast-enhanced
DCE-MRI: dynamic contrast-enhanced magnetic resonance imaging
EES: extravascular-extracellular space
FSL: FMRIB Software Library
GM: gray matter
GRE: gradient-recalled echo
IR: inversion recovery
MRI: magnetic resonance imaging
MTT: mean transit time
NIfTI: Neuroimaging Informatics Technology Initiative (file format)
rICA: right internal carotid artery
ROI: region of interest
SPGR: spoiled gradient recalled echo
SSS: superior sagittal sinus
TE: echo time
TR: repetition time
U-Net: U-shaped convolutional network
VFA: variable flip angle
VOF: venous output function
WM: white matter

## 6 Code and Data Availability

The pipeline is freely available via the *p*-Brain repository on GitHub, complete with detailed installation instructions and thorough documentation. A macOS desktop application for local project organisation and job execution (no cloud upload, data remains on-device) is available at github.com/edtireli/p-brain-platform. Trained CNN weights for slice selection and ROI segmentation are hosted at Zenodo (Tireli, 2025).

## 7 Acknowledgements

The authors thank Antonis Asiminas for valuable discussions and encouragement during the preparation of this work.

## Appendix

### Voxelwise validation against a reference MATLAB implementation

Additional voxelwise comparisons against the reference MATLAB program for perfusion map outputs and for the *T*_1_/*M*_0_ fitting outputs (representative slice 5).

#### Additional cohort-level atlas projections

Additional cohort-mean atlas projections corresponding to the Results section on cohort-level atlas projection.

#### Additional parcel-level maps

Additional parcel-level outputs corresponding to the Results section on regional and parcellated organization.

## Footnotes

1 In this case, segmentation was used from FastSurfer, but in principle any segmentation software may work

2 The boundary is derived as the erosion of the overlap between the dilated gray- and white-matter masks.

